# An anaerobic cell-free protein synthesis formulation for the *in vitro* study of oxygen sensitive proteins

**DOI:** 10.64898/2026.11.09.765404

**Authors:** Janelle D. Arnold, Bo Zhang, Monique N. Milan Lau, Michelle A. O’Malley, Carolyn E. Mills

## Abstract

Anaerobic microbes harbor unique enzymes and metabolic pathways with potential biotechnological applications, yet these microbes remain difficult to study and engineer. Research is limited by the labor-intensive workflows required to maintain anoxic environments and the lack of genetic tools available for non-model anaerobes. Here, we present an anaerobic cell-free protein synthesis (CFPS) platform for the *in vitro* study of oxygen-sensitive enzymes. This study establishes two methods for conducting anaerobic CFPS, achieving an anoxic environment by (1) sparging with inert gas or (2) lyophilizing to remove oxygenated water. We show the functionality of our anaerobic CFPS formulation by producing oxygen-independent fluorescent protein reporters and a [4Fe-4S] cluster pyruvate:ferredoxin oxidoreductase (PFOR) with a specific activity of 5.2 ± 0.3 x 10^-4^ U/µL cell-free extract. This highly modular platform will allow study of diverse oxygen-sensitive proteins and requires only minor modifications to commonly used CFPS workflows.

**Graphical Abstract (created with Biorender):** 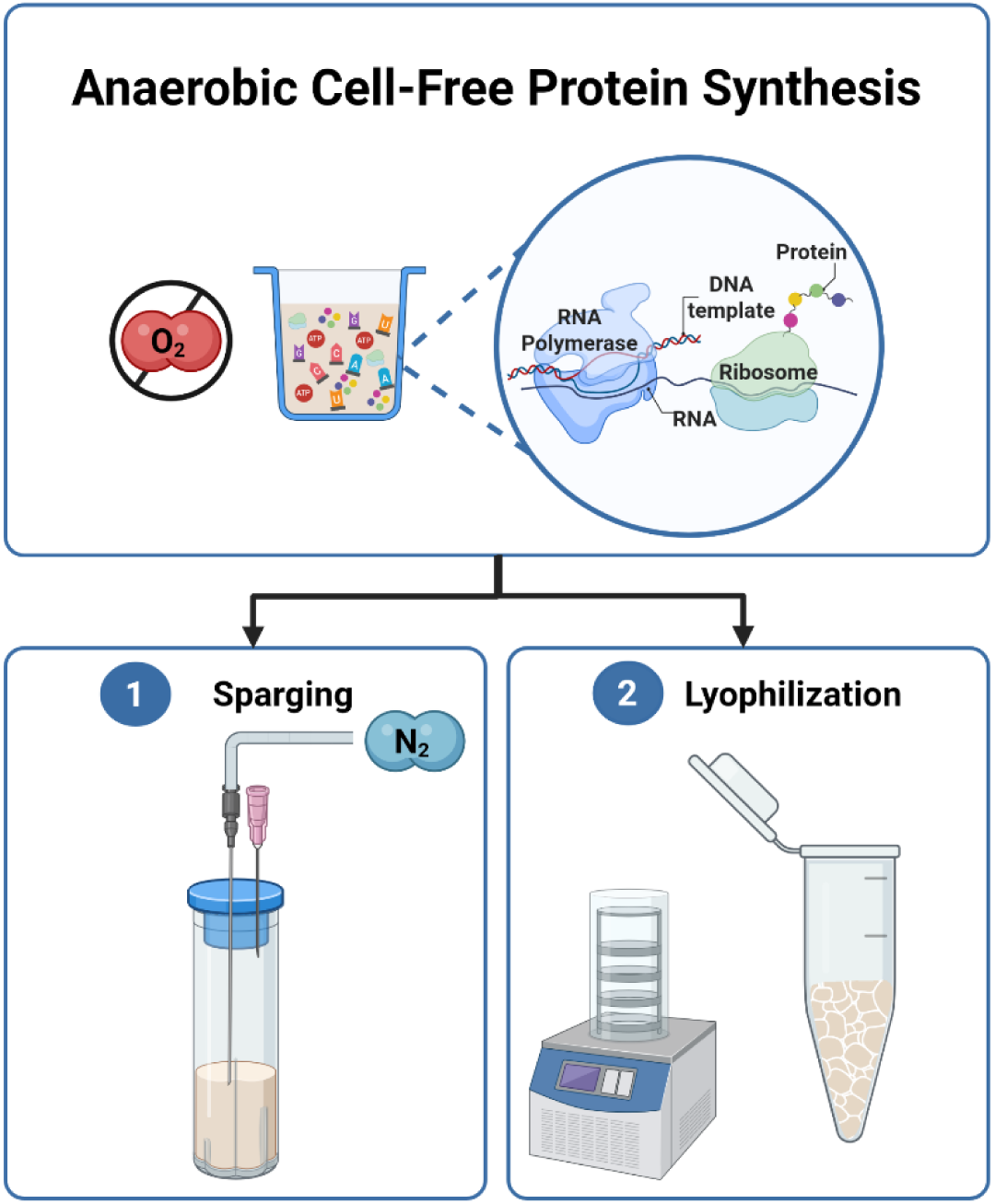

## Introduction

Anaerobic environments, from the human gut to deep ocean geothermal vents, are home to a wide array of microbes that have evolved to thrive in harsh environments with unique enzymes and metabolic pathways adapted to oxygen limitation, steep redox gradients, and other environmental constraints.^1,2^ These features of anaerobes underlie their utility in industrial wastewater treatment, biofuel & biobased chemical production, and specialized fermentation processes.^3–6^ Yet the metabolism of many anaerobic microbes remains understudied, as establishing gene-to-function relationships for such microbes is challenging.^7^ This is due to two major barriers. First, anaerobic microbes as well as essential intracellular enzymes and co-factors are oxygen-sensitive, often requiring the use of anaerobic glove boxes and serum bottles for their study.^1,8^ Second, many anaerobes lack the robust genetic tools that are necessary for knockout studies and the engineering of new functions.^7,9^ Here, we established a platform for the production and study of oxygen sensitive proteins *in vitro* to allow exploration of gene-to-function relationships for anaerobic microbes requiring minimal glovebox use.

Toward characterizing novel proteins and pathways, cell-free protein synthesis (CFPS) systems are an attractive tool for mapping gene-to-function relationships, as they allow *in vitro* transcription and translation of any gene of interest. Beyond simple protein production, CFPS systems offer several unique features that have led to their widespread use in synthetic biology. From a practical standpoint, CFPS produces proteins in an open reaction environment, outside of a living cell. This permits the production of proteins and metabolites that would otherwise be toxic to the cell and streamlines experiments testing combinations of enzymes and/or metabolites that might otherwise require time-consuming sequential transformations – even in environmentally controlled conditions. For more fundamental studies, lysate-based cell-free systems allow reconstitution of the cellular environment without the barrier of cell membranes and cell walls, bridging the disconnect *between in vivo* and traditional *in vitro* experiments.^10^ In this vein, researchers have leveraged cell-free platforms and high-throughput workflows to test the compatibility of genetic circuitry with various model organisms, including *Escherichia coli*, *Saccharomyces cerevisiae*, and HeLa cells, using these CFPS systems as “biomolecular breadboards”.^11–14^ Realizing an accessible anaerobic CFPS system could similarly benefit the study and engineering of anaerobic enzymes and processes.

Prior efforts to produce oxygen-sensitive proteins via CFPS suggest the feasibility and utility of these systems.^15–17^ However, the requirements of anaerobic growth, defined growth media, and housing the equipment required for cell lysis and extract preparation within a glovebox makes such approaches difficult to adopt in laboratories without extensive anaerobic infrastructure. Collectively this prevents the broad application and adoption of anaerobic CFPS platforms. To circumvent some of these challenges, researchers from the Fujishima group produced oxygen-sensitive [4Fe-4S] cluster proteins aerobically. Using the PURE system supplemented with a reconstituted SUF pathway and oxygen scavenging cascade, they created a defined one-pot system for producing oxygen sensitive enzymes at the benchtop.^17^ However, this approach requires the use of multiple purified enzymes and bypasses transcription, using mRNA as a starting material for protein synthesis, making this a time and effort intensive approach for larger scale studies. Other studies have attempted to control the presence of oxygen in CFPS reactions by either passive degassing in an anaerobic glove box^18^ or a methodical tube-in-tube design^19^ to control the passage of oxygen into a cell-free extract. These approaches support the feasibility of anaerobic CFPS but are not widely adoptable for those interested in doing CFPS anaerobically.

In this study, we developed two methods for anaerobic CFPS that simplify the production of oxygen-sensitive proteins. To maximize the accessibility of this technology, we focus on a widely used lysate-based CFPS formulation that can be prepared aerobically using established protocols up until the point of *in vitro* protein synthesis.

Specifically, we use lysates from BL21(DE3) Star *E. coli* with the PANOx-SP energy regeneration system^20,21^. We establish two methods for preparing oxygen-free CFPS reactions—one that uses sparging of inert gases to remove oxygen, and a second that lyophilizes reactions and rehydrates with anoxic water. Using sfGFP and two oxygen-independent fluorescent reporters, we demonstrate robust anaerobic protein production using both methods. Finally, we show that our system can be used to produce a mature [4Fe-4S] cluster containing protein, pyruvate:ferredoxin oxidoreductase (PFOR) from the anaerobic bacterium *Prevotella brevis* GA33—an oxygen-sensitive enzyme involved in anaerobic central metabolism^22^.

## Results and Discussion

### Sparging cell-free extracts with nitrogen gas is sufficient to achieve a fully anaerobic system for in vitro transcription and translation

We began by establishing a reliable readout for protein expression under anoxic conditions to test the productivity of our anaerobic CFPS formulations. Commonly, cell-free formulations are optimized using superfolder GFP (sfGFP), where fluorescence serves as a proxy for protein yield and higher fluorescence indicates higher concentrations of the protein product of interest.^23,24^ However, sfGFP requires oxygen for chromophore formation, precluding direct measurement of protein production under anaerobic conditions via fluorescence.^25,26^ To circumvent this issue, we quantified anaerobic CFPS yields using two oxygen-independent fluorescent reporter proteins: iRFP702,^27^ a near-infrared fluorescent protein validated anaerobically *in vivo*,^28^ and mFAP2a,^29^ a *de novo*-designed fluorescent protein (Table 1). Both proteins fluoresce upon binding to a cofactor—biliverdin for iRFP702 and DFHBI for mFAP2a—that can be added directly to CFPS reactions. In addition, we use sfGFP in this study as an *in situ* oxygen indicator to determine if anoxic conditions were maintained during protein synthesis.

**Table 1.**
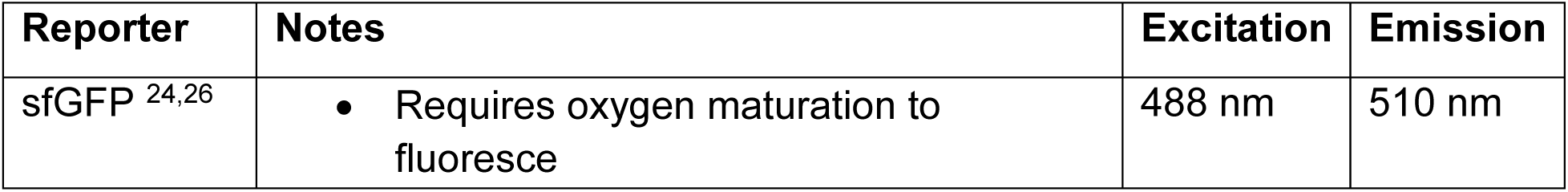
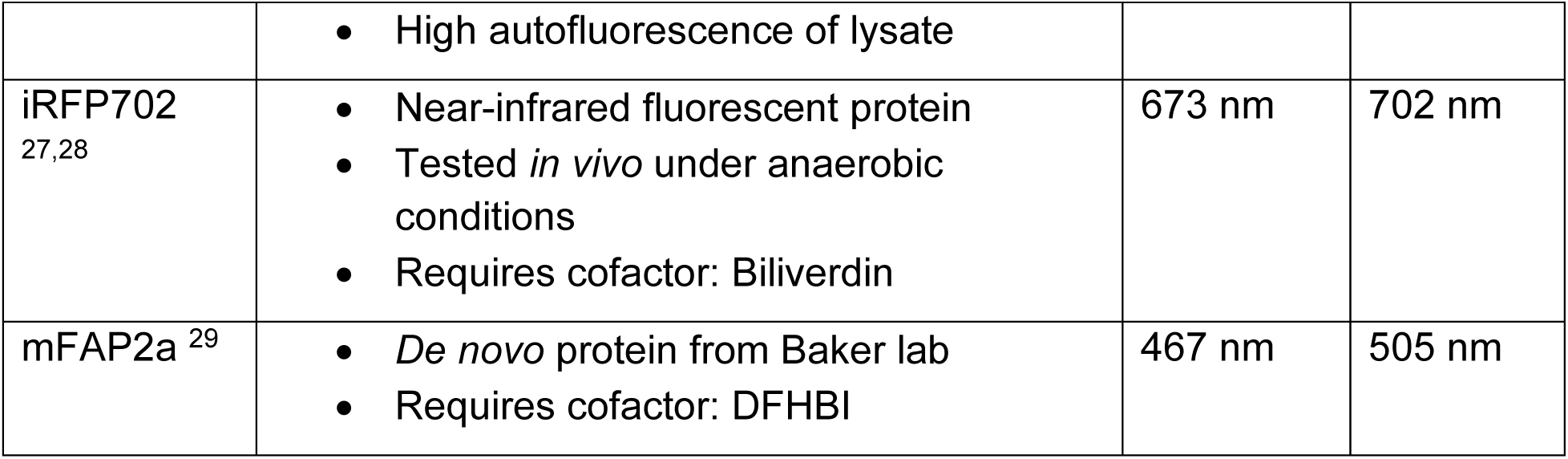
Summary of fluorescent reporter proteins used in this study.

| Reporter | Notes | Excitation | Emission |
|---|---|---|---|
| sfGFP<sup>24,26</sup> | • Requires oxygen maturation to fluoresce<br>• High autofluorescence of lysate | 488 nm | 510 nm |
| iRFP702<sup>27,28</sup> | • Near-infrared fluorescent protein<br>• Tested *in vivo* under anaerobic conditions<br>• Requires cofactor: Biliverdin | 673 nm | 702 nm |
| mFAP2a<sup>29</sup> | • *De novo* protein from Baker lab<br>• Requires cofactor: DFHBI | 467 nm | 505 nm |

Our primary goal was to establish a “treatment” that removes oxygen from a CFPS reaction after its preparation under aerobic conditions. Treatment options were limited to those that have minimal impact on the structure and activity of the biomolecules required for *in vitro* transcription and translation. Drawing inspiration from anaerobic microbial culture techniques, we began by exploring sparging methods commonly used in anaerobic media preparation as a route to removing oxygen from our cell-free system (Figure 1). First, we mixed CFPS reagents under constant flow of nitrogen gas in a glass cuvette fit to seal with a butyl rubber stopper (Figure 2A). Once all reaction components were mixed, we sealed the cuvette with a stopper and created a gas inlet/outlet with a 2-needle system (Figure 2B). Nitrogen gas was gently bubbled into the cell-free extract effectively displacing dissolved oxygen with nitrogen, an inert gas.

**Figure 1.**
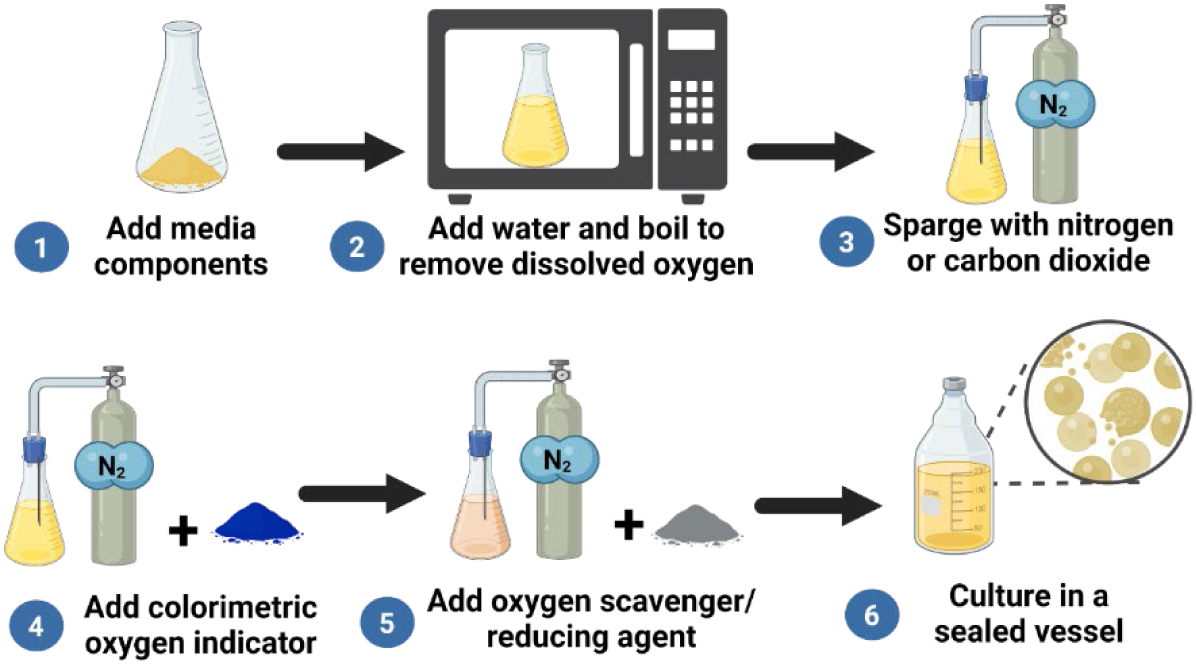
Overview of methods for preparing media to culture anaerobic microbes. Figure generated with Biorender.

**Figure 2.**
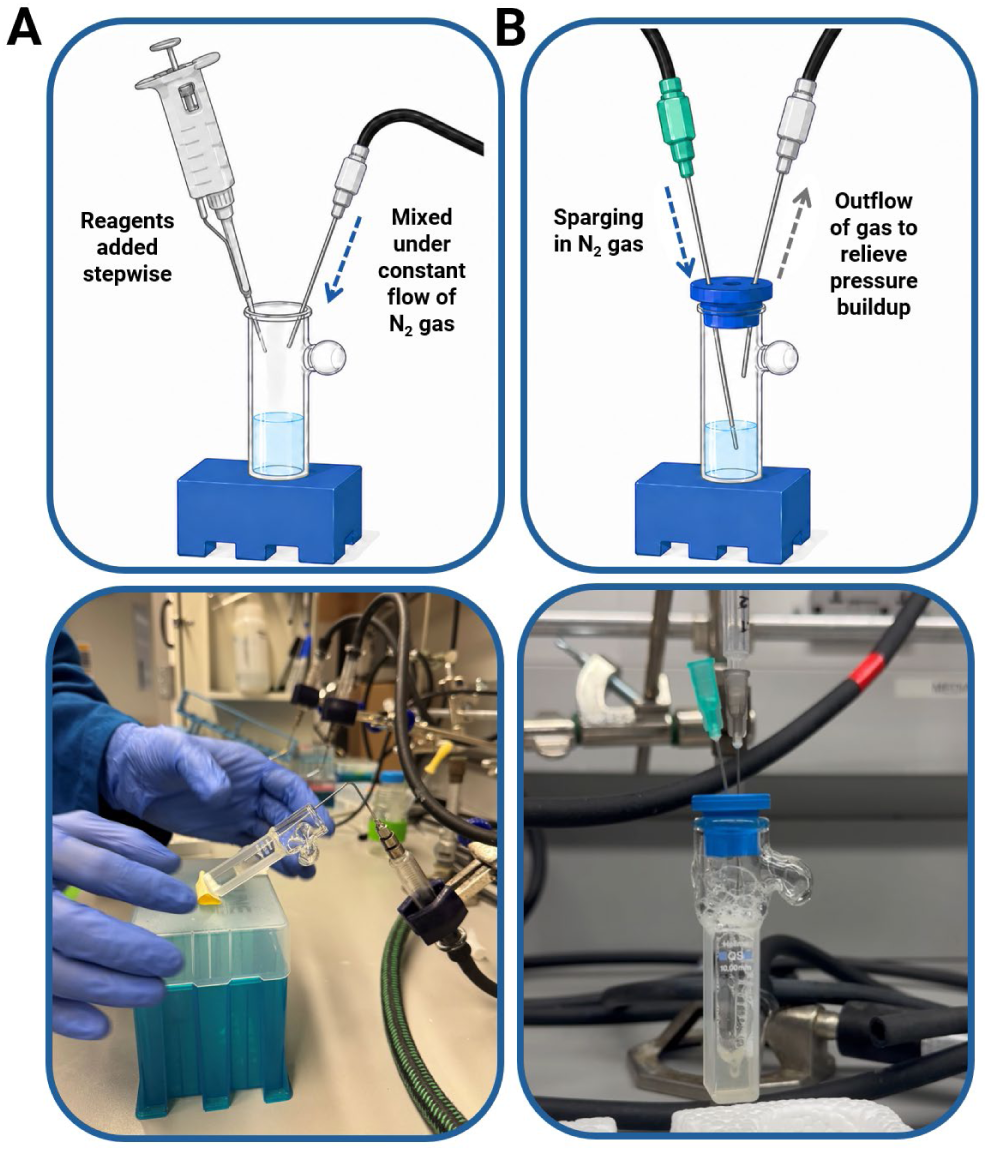
Cell-free extracts were purged of oxygen by sparging in nitrogen gas. (A) Oxygenated cell-free reaction components were mixed under the constant flow of nitrogen gas in a glass cuvette. (B) After all reaction components were mixed, the cuvette was sealed with a rubber septum. Using two needles, we created an inlet and outlet to bubble in nitrogen without allowing pressure to build in the sealed cuvette. This master mix was then transported to the anaerobic chamber where it was aliquoted into individual reactions. Top panels of this figure were generated with ChatGPT 5.6-Sol.

We found that sparging with nitrogen gas is sufficient to remove oxygen from solution and that CFPS still proceeds under anaerobic conditions. sfGFP fluorescence was used as an oxygen indicator to determine if we achieved an anoxic environment and the fluorescence produced by our two oxygen-independent reporters (iRFP702, mFAP2a) as a proxy for protein yield. Specifically, if an anaerobic environment is maintained during protein synthesis, sfGFP will be produced, but will not fluoresce until exposed to oxygen and post-translational oxidation allows GFP chromophore formation to proceed. In contrast, iRFP702 and mFAP2a fluorescence should remain the same before and after oxygen exposure as their fluorescence is independent of oxygen. After allowing the CFPS reaction to proceed for 24 hours, we measured fluorescence of all reporters while the plate was still sealed with an optically clear, gas impermeable seal (Figure 3). We then removed the seal, halted translation with the addition of chloramphenicol, and monitored sfGFP fluorescence for 90 minutes. These data indicated that sparging the reaction mixture prior to DNA addition was sufficient for achieving an anoxic environment and that protein synthesis proceeded in these reactions in the absence of oxygen—after the 24-hour incubation but before oxygen exposure, no fluorescence was observed in reactions producing sfGFP while both iRFP702 and mFAP2a reactions produced a fluorescent signal (Figure 4A). After oxygen exposure, fluorescence in iRFP702 and mFAP2a CFPS reactions did not change (*p_iRFP702_* = 0.66, *p_mFAP2a_* = 0.55, 2-tailed *t*-test), whereas the observed fluorescence in the sfGFP reaction increased substantially (*p* = 0.004, Figure 4A).

**Figure 3.**
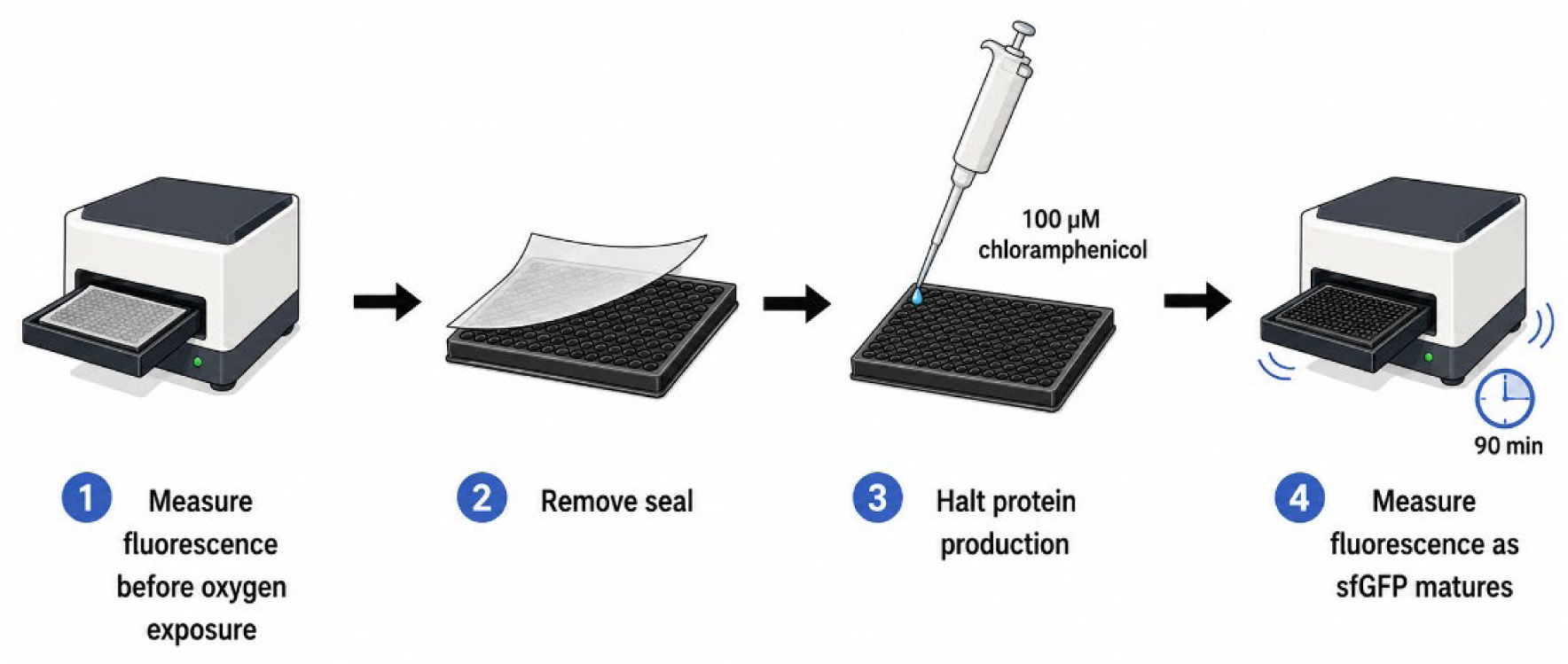
sfGFP serves as a proxy for protein yields and oxygen exposure in anaerobic cell-free reactions. All anaerobic CFPS experiments in this study were conducted in a 384-well microplate sealed with a gas-impermeable clear adhesive. Reactions were prepared and sealed in the anaerobic chamber, then removed and incubated for 24 hours at 30°C. Fluorescence measurements were taken before and after removing the adhesive seal, marking the transition from an anoxic to aerobic environment post-protein synthesis. sfGFP fluorescence was measured over a 90-minute time course with orbital shaking (180 rpm) and measurements taken every 60 seconds. Since sfGFP requires the presence of oxygen to fluoresce, the change in fluorescence before and after oxygen exposure would indicate that we had achieved an anoxic environment for protein synthesis. Chloramphenicol was added to halt translation and ensure the change in fluorescence of sfGFP was due to oxygen binding to protein produced anaerobically and not new protein production once exposed to oxygen. This schematic was generated with ChatGPT 5.6-Sol.

**Figure 4.**
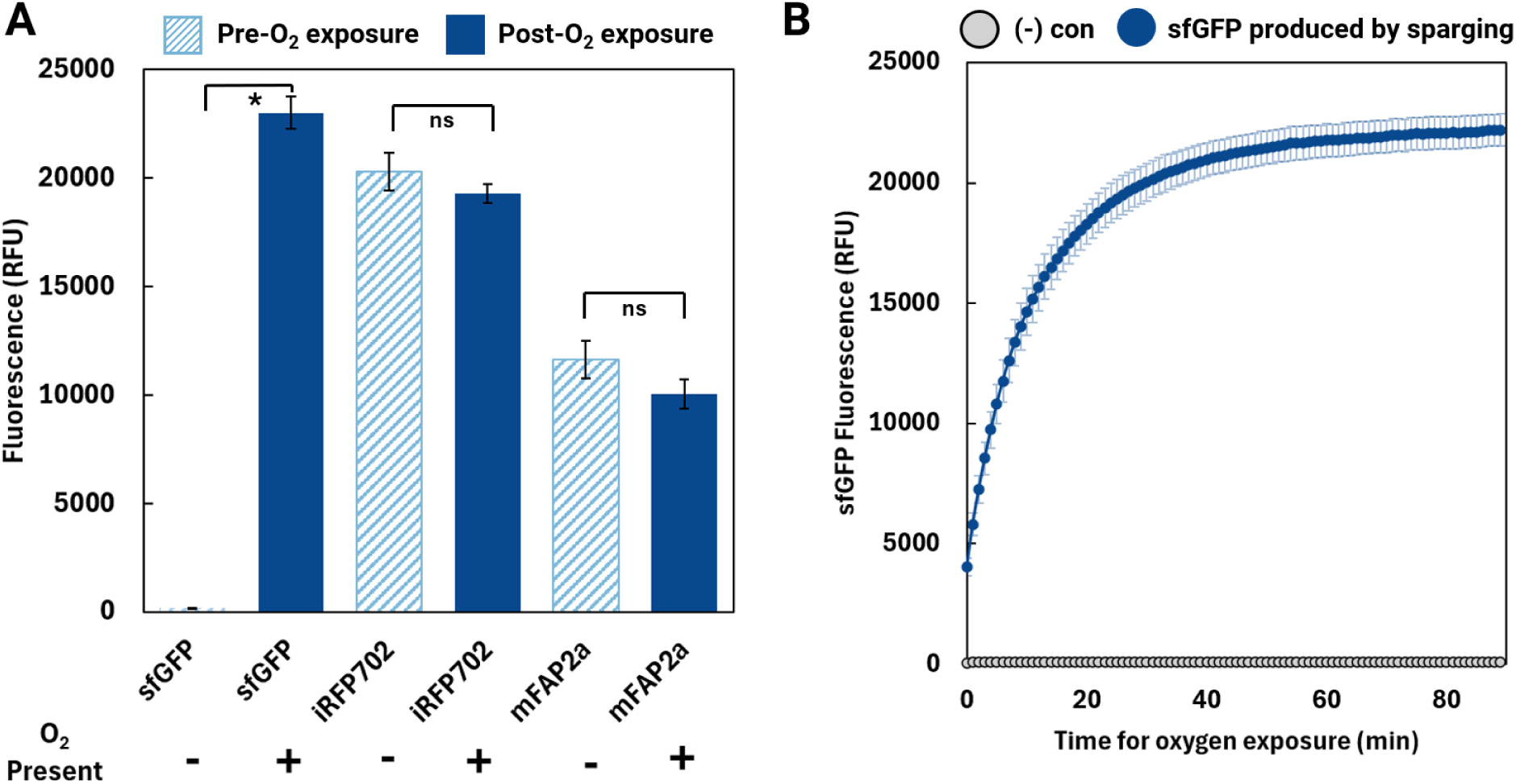
Anaerobic CFPS performed via sparging with nitrogen gas. (A) Relative fluorescence of 3 fluorescent reporter proteins above background indicate protein was produced anaerobically. After incubating for 24 hours at 30°C in sealed 384-well plates, fluorescence measurements were taken. Then, the seal was removed, translation was halted with the addition of 100uM chloramphenicol, and fluorescence was measured again after 90 minutes of exposure to oxygen. This data represents fluorescence before and after oxygen exposure with sfGFP fluorescence maturing after exposure and iRFP702 and mFAP2a fluorescence remaining constant. (B) sfGFP fluorescence increasing rapidly during oxygen exposure represents protein maturation, rather than new protein production. sfGFP can be produced anaerobically, however, it requires oxygen to bind to mature and fluoresce. This mechanism allowed us to use sfGFP as an oxygen indictor in our system with no fluorescence indicating an anaerobic environment and the rapid increase in fluorescence indicating the addition of oxygen to our system. (A-B) Error bars represent ±1 standard error of the mean for n=4 sample size.

Interestingly, this setup also allowed us to track sfGFP chromophore maturation as a function of time after oxygen exposure (Figure 4B). Previous studies have tracked chromophore maturation of *de novo* GFP variants using the PURE CFPS system coupled with an oxygen scavenging cascade that used glucose, glucose oxidase, and catalase.^25^ Our method produced similar kinetic data for sfGFP chromophore maturation while avoiding the need to externally supplement high-cost enzymes, suggesting the potential utility of crude extract-based CFPS systems in the kinetic study of fluorescent protein chromophore maturation.

In early experiments where chloramphenicol was not added to halt translation we observed both sfGFP chromophore maturation immediately after oxygen exposure and, after a delay, the production of new sfGFP (Figure S1). We did not expect oxygen exposure to facilitate new protein production based on prior results performing aerobic CFPS using this same formulation—under aerobic conditions, protein production in this same CFPS formulation begins to plateau after 10 hours and ATP concentrations reach zero after 24 hours.^23^ Thus, we expected our anaerobic CFPS reaction to be similarly depleted, as these reactions had already been incubated for 24 hours at 30 °C; however, this was not the case. The role of oxygen in controlling protein yields for this formulation is discussed in more depth in later sections.

In early iterations of this work, we also explored the use of strong reducing agents as oxygen scavengers and resazurin as an oxygen indicator; however, we found that reducing agents were unnecessary to achieve an anoxic environment and ultimately lowered protein yields (Figure S2). Resazurin is a chemical redox indicator commonly added to anaerobic media; when reduced to resorufin, this serves as a colorimetric (A570 nm) and fluorogenic (571 nm excitation/ 584 nm emission) oxygen indicator.^30^ After producing our fluorescent reporter proteins anaerobically, we determined if the reaction was anoxic by removing the seal and exposing it to oxygen (Figure 3). We monitored the change in resorufin fluorescence as a proxy for oxygen dissolution because it is a more sensitive measurement than absorbance and is agnostic to aggregation that could occur during CFPS reactions (Figure 5). In these experiments, resazurin was always coupled with cysteine-HCl, to ensure it was reduced to resorufin.

**Figure 5.**
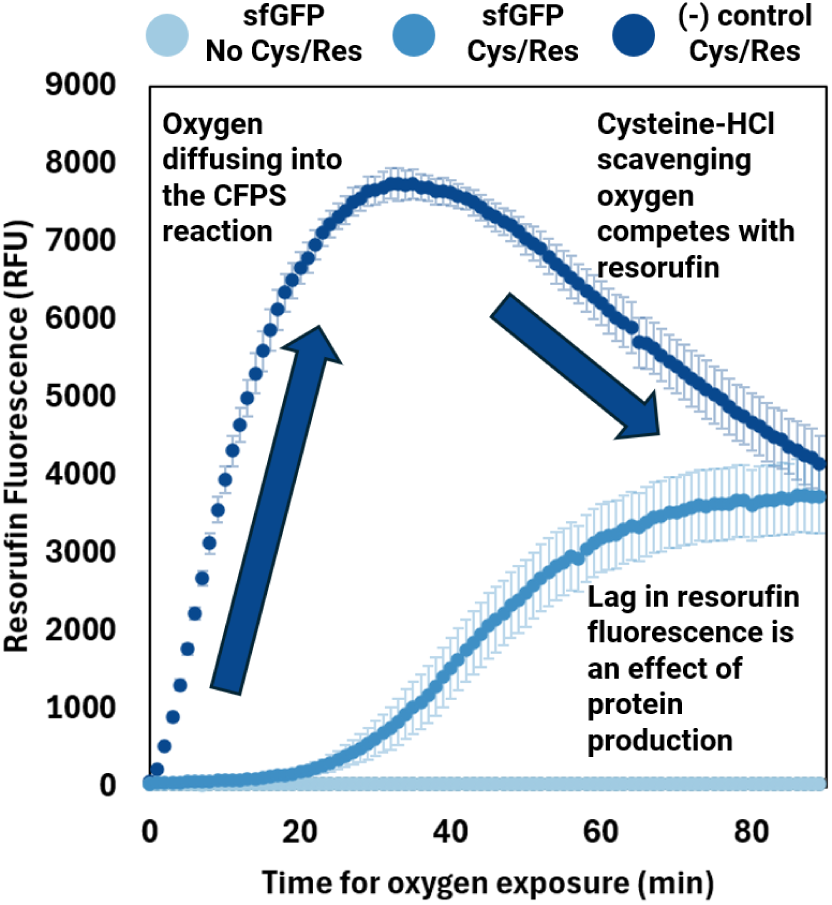
Resorufin serves as an oxygen indicator in anaerobic cell-free reactions. Resazurin is first reduced to resorufin (by cysteine-HCl in this experiment) before serving as a fluorogenic oxygen indicator. Resorufin fluorescence (571 nm excitation/ 584 nm emission) increases as oxygen diffuses into the previously anoxic CFPS reaction. Error bars represent the standard error of the mean (n=4).

The negative control containing resazurin/cysteine-HCl and omitting DNA showed a rapid rise in fluorescence as resorufin reacted with oxygen after exposure. This indicated that our reactions were free of oxygen during protein synthesis, in agreement with sfGFP data. After 30 minutes, resorufin fluorescence reached a peak before decreasing, most likely driven by cysteine-HCl scavenging oxygen. The control expressing sfGFP but omitting resazurin/cysteine-HCl did not show any fluorescence during the 90-minute time course, indicating that sfGFP did not contribute to the observed fluorescence in this channel. The experimental condition in which sfGFP was expressed anaerobically, and resazurin/cysteine-HCl were present, did not immediately fluoresce upon exposure to oxygen. Instead, a lag-phase was observed for approximately 20 minutes before oxygen exposure produced an increase in resorufin fluorescence. This trend appeared across all fluorescent reporter proteins (Figure S3) and was not specific to sfGFP, suggesting that the lag in observed resorufin fluorescence upon oxygen exposure is associated with CFPS reactions that have produced protein. While interesting, these differences in behavior of resorufin fluorescence are beyond the scope of this study. We instead chose to use sfGFP as an *in-situ* oxygen indicator for subsequent experiments.

### Lyophilization streamlines batch-prepping of cell-free extracts for anaerobic in vitro transcription and translation

Lyophilization is a well-established technique that enables long-term storage and transport of CFPS reactions for use in on-site vaccine manufacturing, diagnostics, and water quality sensing^31–34^. Here, we explored lyophilization as a route to streamline the setup of anaerobic CFPS reactions (Figure 6). Specifically, we lyophilized pre-mixed cell-free reactions to remove oxygenated water through sublimination while preserving the activity of reagents required for transcription and translation. Freeze-dried pellets were then rehydrated in an anaerobic chamber with anoxic water to produce an anaerobic CFPS reaction mixture. This rehydrated master mix was then aliquoted into individual reactions, after which DNA and protein-specific co-factors were added. Plates were then sealed and incubated to allow protein synthesis to proceed.

**Figure 6.**
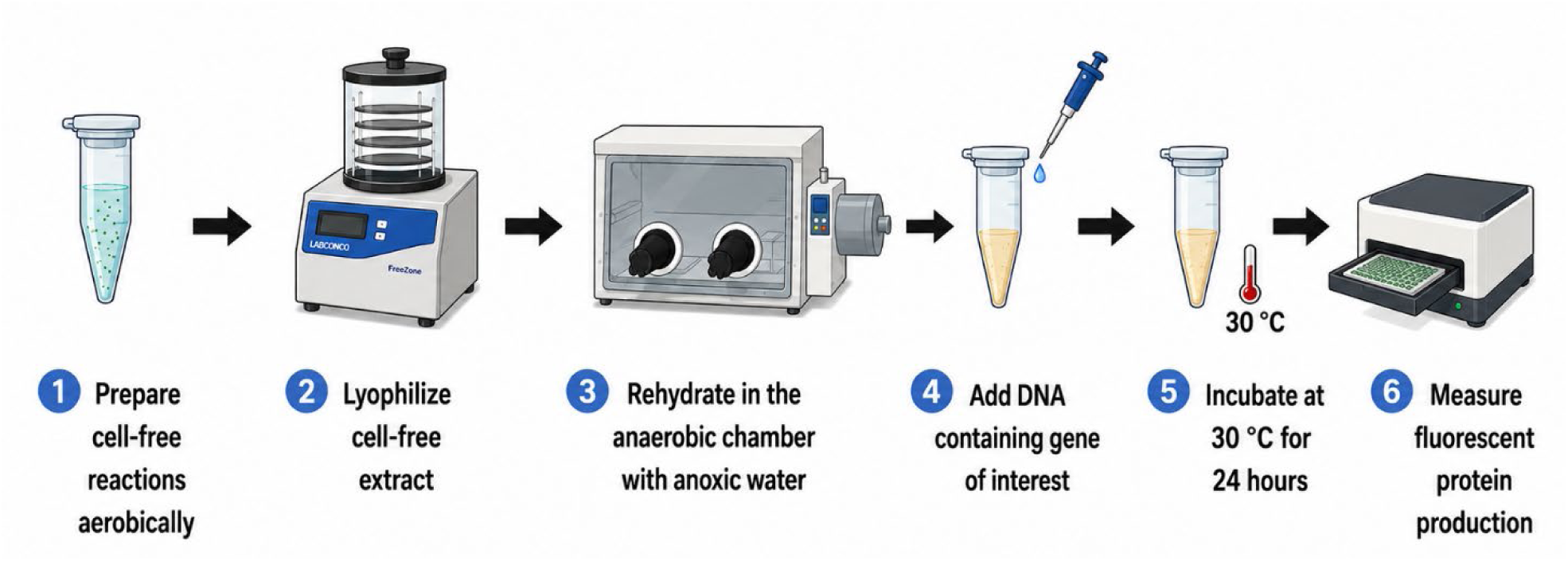
Oxygen was be removed from cell-free extracts by lyophilizing samples and rehydrating them with anoxic water. Cell-free extracts were mixed and prepared aerobically before being lyophilized for 16 hours. Lyophilization removed oxygenated water from the samples without disturbing the integrity of proteins and small molecules in our reactions.Lyophilized samples were then rehydrated in the anaerobic chamber with anoxic water (nitrogen flushed) and aliquoted into individual reactions. This schematic was generated with ChatGPT 5.6-Sol.

We applied this lyophilization/rehydration/anaerobic (LyRA) workflow to our CFPS reactions and achieved anaerobic protein synthesis with similar yields to sparged CFPS reactions for two of our three reporters (Figure 7A). First, we verified that LyRA CFPS reactions were anaerobic using the same sfGFP chromophore maturation experiment we used with sparged CFPS reactions (Figure 7B). Similar to our nitrogen-sparged CFPS reactions, LyRA CFPS reactions producing sfGFP did not fluoresce until the plate was unsealed and exposed to oxygen, indicating we maintained anoxic conditions.

**Figure 7.**
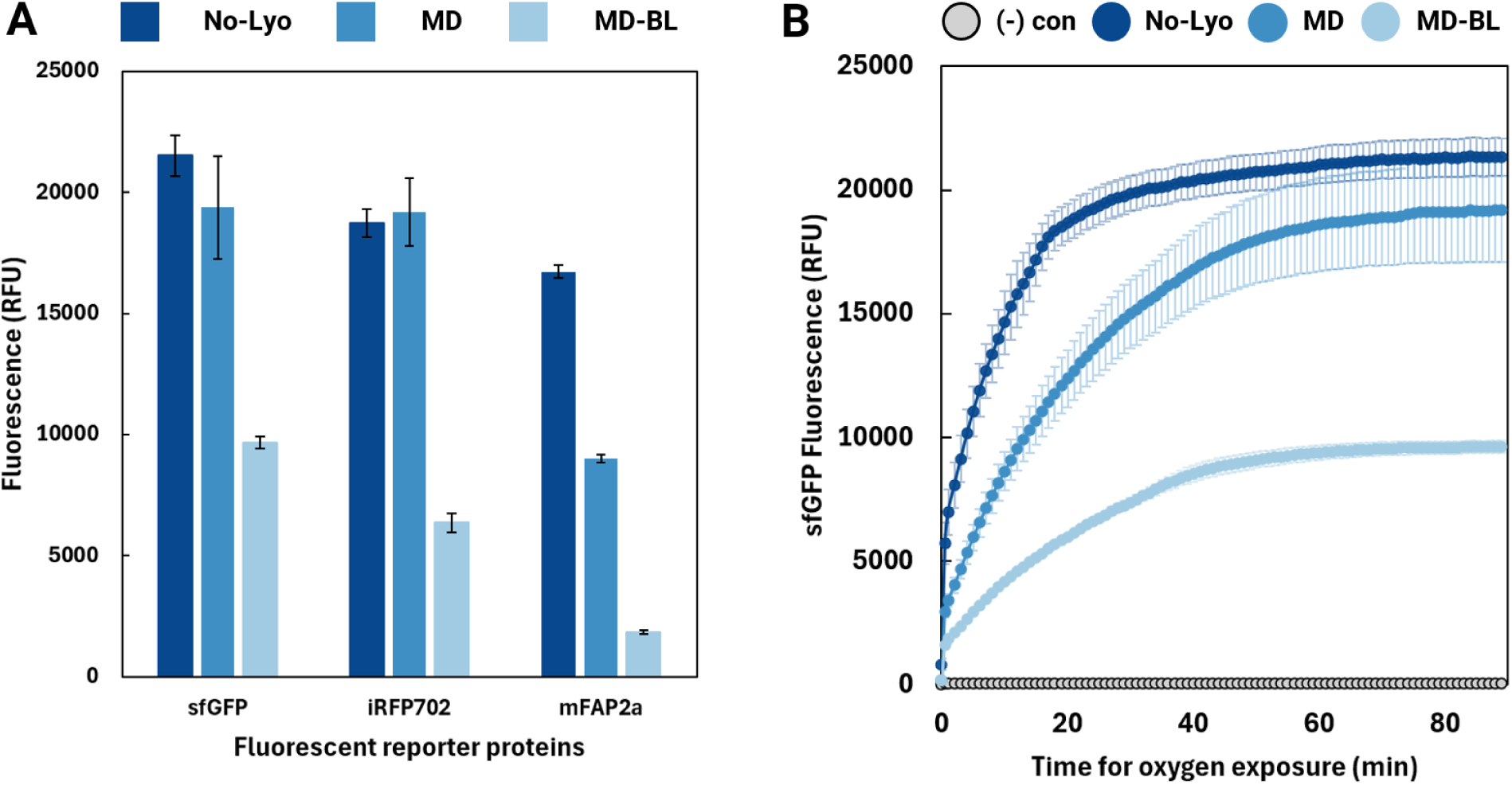
Anaerobic CFPS performed via lyophilization and rehydration with anoxic water (LyRA). We measured relative protein production under anaerobic conditions in 2 different CFPS formulations optimized for lyophilization (maltodextrin and maltodextrin/ β-lactose) including a control with no lyoprotectants. All 3 formulations produced protein as measured by fluorescence above background, with the no lyoprotectant control producing the highest yields. After incubating CFPS reactions for 24 hours at 30 °C in sealed 384-well plates, fluorescence was measured in a plate reader at their respective emissions and excitations. Then, the seal was removed, translation was halted with the addition of 100uM chloramphenicol, and fluorescence was measured again after 90 minutes of exposure to oxygen. This data represents fluorescence before and after oxygen exposure with sfGFP fluorescence maturing after exposure and iRFP702 and mFAP2a fluorescence remaining constant. (B) sfGFP fluorescence increases over time as exposed to oxygen indicating an anoxic environment was achieved during protein synthesis with the lyophilization technique. (A-B) Error bars represent ±1 standard error mean for n=3 sample size.

Comparing reporter yields across sparged and LyRA CFPS reactions, we find that sfGFP and iRFP702 yields were not statistically different across the two different oxygen-removal strategies (*p*_sfGFP_=0.55, *p*_iRFP702_=0.71, 2-tailed *t*-test). In contrast, mFAP2a yields increased approximately 70% in the lyophilized anaerobic CFPS reaction. The difference in yields in sparged versus LyRA CFPS reactions for only one of our three reporters suggests that exploring both deoxygenation strategies may be beneficial when attempting to optimize anaerobic CFPS yields; however, because LyRA CFPS reactions produced similar or higher yields across all our reporters, we concluded that this is a viable approach for facile preparation of anaerobic CFPS reactions.

After establishing LyRA for our baseline CFPS formulation, we quantified the impact of lyoprotectants on final protein yields to determine whether their addition could improve either baseline protein production or stability of lyophilized reactions upon storage. We tested two different sugar additives previously reported as lyoprotectants for CFPS reactions: maltodextrin (MD) and maltodextrin + β-lactose (MDBL). These lyoprotectants were selected because they have been reported across multiple groups to be effective at improving the thermostability of CFPS samples, thereby enabling long-term storage.^32,33^ The no-lyoprotectant control and the MD formulation produced similar concentrations of protein for both sfGFP and iRFP702 (*p*_sfGFP_=0.64, *p*_iRFP702_=0.87, 2-tailed *t*-test), which is consistent with prior reports that MD as an additive in the PANOx-SP system does not alter CFPS yields for sfGFP (Figure 7A).^33^ Interestingly, the addition of maltodextrin did decrease the yield of mFAP2a fluorescence by 46 ± 3% compared to the no lyoprotectant control. While a study of how lyoprotectants impact the specific yields of different proteins is beyond the scope of this work, we hypothesize that this could be the result of maltodextrin shielding access to DFHBI, the co-factor required for mFAP2a fluorescence.

From a practical standpoint, the MD LyRA CFPS formulation was easier to reconstitute compared to the no-lyoprotectant control, as it produced fewer bubbles during rehydration. This is expected, as maltodextrin is known to decrease the cake porosity of samples dried during lyophilization, thereby protecting biologics during the lyophilization process and allowing easier reconstitution.^35–37^ Given that the addition of maltodextrin improves processibility while minimally affecting yield, it is a good candidate for experiments where researchers may have limited control over pipetting during rehydration, which may be the case for some anaerobic chambers or automated workflows. Tracking sfGFP chromophore maturation, we see that all lyoprotectant formulations remained anoxic during protein synthesis, with no difference in final yields (*p*=0.64, 2-tailed *t*-test) between the no-lyoprotectant control and the MD formulations (Figure 7B).

In contrast to the no lyoprotectant control and maltodextrin alone, the addition of both maltodextrin and β-lactose consistently decreased LyRA CFPS protein yields for all reporters tested. Specifically, sfGFP, iRFP702, and mFAP2a yields in reactions with MDBL were 55%, 66%, and 89% lower than the no lyoprotectant control, respectively. This is consistent with previous literature showing that most sugar-based lyoprotectants negatively impact CFPS yields for samples rehydrated immediately after lyophilization. Specifically, one study screening the effect of lyoprotectants on sfGFP yields found that 5 out of 6 lyoprotectants, including β-lactose, significantly lowered protein yields when CFPS reactions were rehydrated immediately after lyophilization;^32^ consistent with our results, the only exception to this trend was maltodextrin. In these studies, lyoprotectants are used to extend the shelf life of CFPS reactions stored at room temperature or higher, however, because our interest lies in characterizing gene-to-function relationships for anaerobic microbes, cold storage is readily available, and we instead focus on formulations that maximize overall yield.

Combined, these results indicate that LyRA offers a reproducible and streamlined method for preparing anaerobic cell-free extracts while eliminating the need for time-intensive setup of a sparging apparatus the day of experimentation. Specifically, lyophilization removes variability associated with nitrogen bubbling flow rates, the formation of unwanted foam, and provides more flexibility in the volumes that can be prepared for a CFPS experiment. In the LyRA workflow proposed and tested here, both the lyophilized extracts and anoxic water can be prepared in large batches and stored for long-term use. Anoxic water is shelf-stable in serum bottles sealed with a rubber butyl stopper and crimped, and lyophilized cell-free reactions are stable for multiple months in cold storage conditions (-80 °C and -20 °C).^31^ This LyRA workflow therefore significantly reduces experimental time for oxygen sensitive proteins produced via CFPS, requiring only same-day rehydration and the addition of DNA.

Next, we sought to establish whether long-term storage impacts the yields of our LyRA CFPS reactions. To this end, we prepared a large batch of lyophilized CFPS reaction mixes with and without lyoprotectants, stored them at different temperatures (room temperature, -20 °C, -80 °C) and rehydrated aliquots once a week under aerobic and anaerobic conditions for 4 weeks. We rehydrated aliquots both aerobically and anaerobically to determine whether storage differentially impacts *in vitro* protein synthesis carried out in the presence and absence of oxygen. We used sfGFP as the reporter for aerobic protein synthesis yields and iRFP702 as the reporter for anaerobic protein synthesis yields.

We found that long-term cold storage (-20 °C and -80 °C) of lyophilized CFPS reactions does not lead to a significant loss in final protein yield upon rehydration in both aerobic and anaerobic CFPS reactions. After 4 weeks of storage at -20 °C and -80 °C, CFPS reactions rehydrated aerobically maintained comparable yields to those rehydrated immediately after lyophilization, regardless of lyoprotectant addition or the identity of the lyoprotectant (Figure S4). However, 4 weeks of room temperature storage resulted in a 58-92% decrease in the yields of CFPS reactions rehydrated aerobically, depending on the lyoprotectant added. This is consistent with prior reports that long-term storage at elevated temperatures leads to significant losses in the final yield of CFPS reactions— despite maintaining higher activity than the no lyoprotectant control, yields still decrease for both MD and MDBL after 4 weeks with room temperature storage.^33^

Similar to aerobically rehydrated CFPS reactions, LyRA CFPS yields for the no lyoprotectant and MD formulations were not significantly different between week 0 and week 4 for reactions stored at -20 °C and -80 °C (*p_-20 No Lyo_*=0.052, *p_-20 MD_*=0.32, *p_-80 No Lyo_*=0.052, *p_-80 MD_*=0.46, 2-tailed *t*-test; Figure 8). The MDBL formulation, however, did suffer yield losses upon storage, with a 27 ± 2 % decrease in yield when stored at -20°C and 37 ± 4 % when stored at -80 °C. For LyRA CFPS samples without lyoprotectants, storage at room temperature led to an immediate drop-in activity, retaining only 6% of the initial yield by week 2. In contrast, the MD formulation LyRA yields were not significantly different between week 0 and week 4 when stored at room temperature (*p_RT MD_*=0.066, 2-tailed *t*-test). In summary, cold storage consistently improved the retention of activity in LyRA CFPS reactions, allowing us to move forward without the use of lyoprotectants and long-term storage at -20 °C when preparing large batches of reaction mixtures.

**Figure 8.**
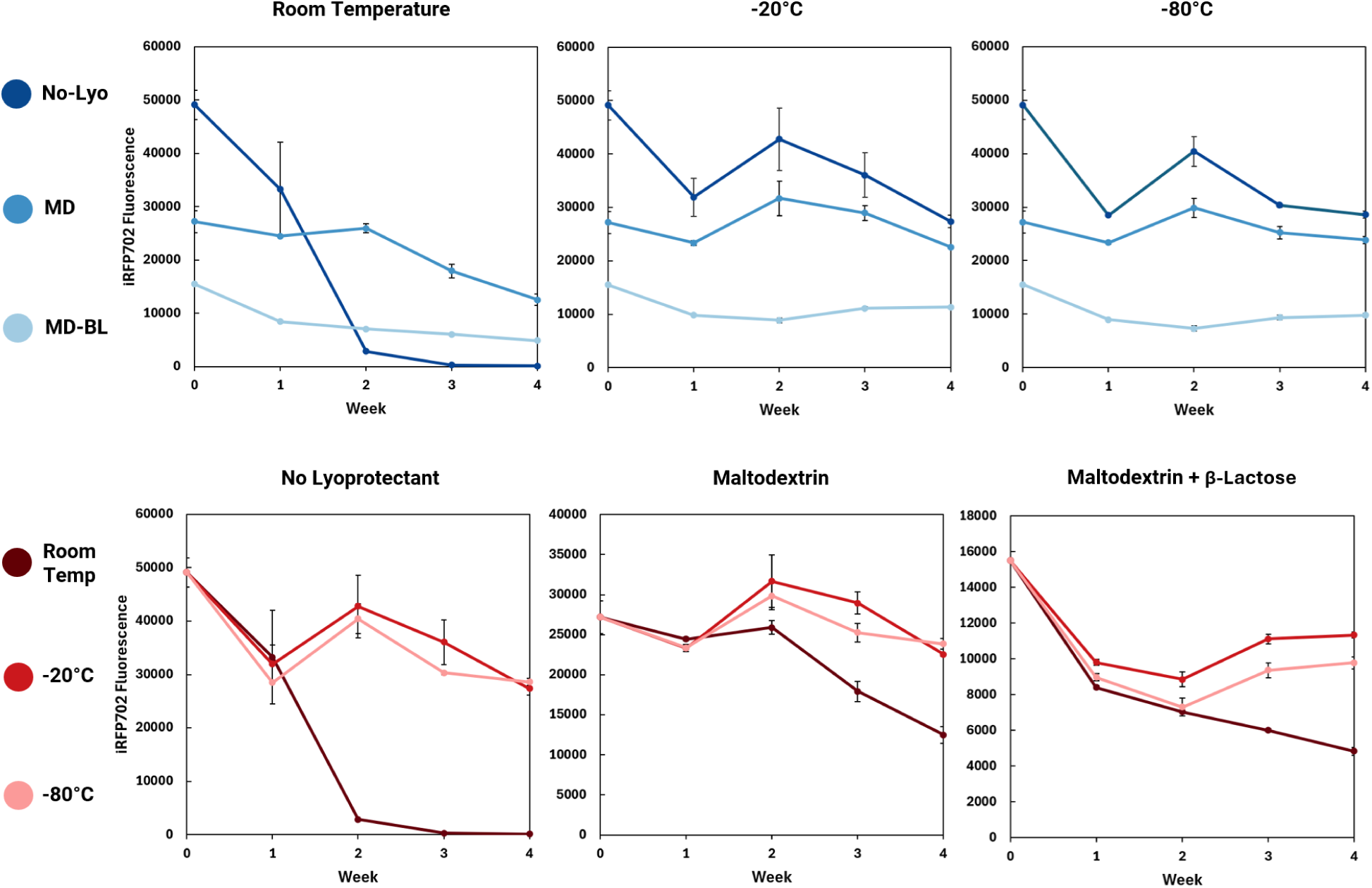
Optimization of formulation and storage conditions for LyRA CFPS reactions. LyRA CFPS reactions were lyophilized in a large batch of aliquots and stored at room temperature, -20 °C, or -80 °C for up to 4 weeks. Each week, an aliquot was removed and rehydrated with anoxic water to determine the level of activity retained after long-term storage at that temperature.

### Anaerobic CFPS yields less protein compared to aerobic reactions

This study used the PANOx-SP CFPS formulation, which relies on PEP as the primary energy source for ATP regeneration. Specifically, the transformation of PEP into pyruvate by pyruvate kinase converts one ADP into ATP per PEP molecule processed— a process that does not rely on oxygen to proceed. NAD+ and CoA supplementation facilitates the regeneration of additional ATP via a series of similarly oxygen-independent reactions that convert pyruvate to acetate.^38^ Combined, these features of PANOx-SP suggest that aerobic and anaerobic CFPS yields should be similar. However, while these primary modes of ATP regeneration in PANOx-SP CFPS formulations are oxygen-independent, previous work established that protein yields can exhibit oxygen dependence for any cell-free formulations that contain native cytoplasmic energy regeneration system found in crude cell extracts.^39^ Further, recent campaigns to improve protein yields in CFPS have shown that supplementing oxygen to crude-extract CFPS reactions can substantially increase yield.^23^

Given these observations, we sought to quantify changes in protein yield between our anaerobic and aerobic CFPS reactions. To quantify these changes, we first sparged a pre-mixed cell-free reaction with nitrogen gas, as in our sparging experiments above (Figure 9A). We then split this sparged CFPS reaction mix into two separate batches— one to be used for anaerobic protein production and one to be used for aerobic protein production. Importantly, sparging the entire reaction mix with nitrogen ensured that any evaporation or protein damage related to sparging was consistent between both conditions such that any differences in yield could be directly attributed to oxygen availability. After splitting the CFPS reaction mixture, DNA and protein-specific cofactors were added to the anaerobic reaction under a N_2_/H_2_ headspace in an anaerobic chamber and to the aerobic reaction at the benchtop, where it was exposed to oxygen. The anaerobic and aerobic samples were both sealed to prevent evaporation during incubation; however, the aerobic sample was sealed with a gas-permeable seal to allow oxygen transport into the microwell plate.

**Figure 9.**
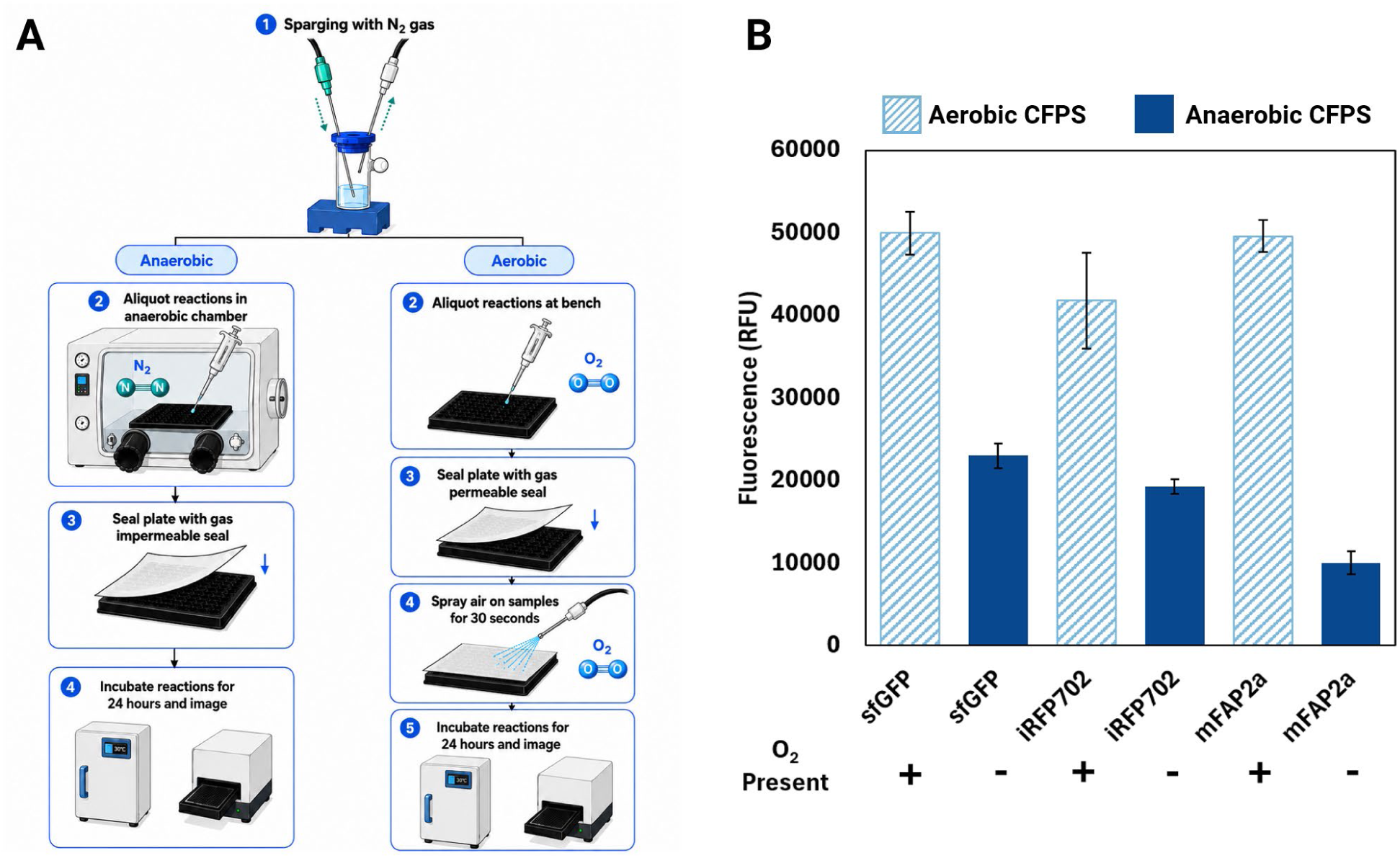
Comparison of protein production using the PANOX-SP energy regeneration CFPS system under anaerobic and aerobic conditions. (A) Experimental workflow of preparing anaerobic and aerobic cell-free extracts. Both anaerobic and aerobic samples were degassed with nitrogen to remove oxygen and account for potential evaporation effects during this treatment. Anaerobic samples were then aliquoted into a 384-well plate with a gas-impermeable seal in the anaerobic chamber. Aerobic samples were there aliquoted into a 384-well plate at the benchtop, introducing oxygen through pipetting. Samples were sealed with a gas-permeable seal that allows for the flow of oxygen but reduces evaporation and sprayed with house air for 30 seconds to ensure adequate oxygen exposure. After incubating reactions for 24 hours at 30°C, seals were removed and all reactions were halted with the addition of 100uM chloramphenicol, and fluorescence was measured after 90 minutes of exposure to oxygen. This schematic was generated with ChatGPT 5.6-Sol.(B) Anaerobic CFPS yields significantly less protein compared to aerobic production across all three fluorescent reporters. All fluorescent data presented here is after oxygen exposure. Error bars represent ±1 standard deviation for n=4 sample size.

We observed a 60% decrease in protein yield when comparing anaerobic versus aerobic CFPS reactions, averaged across all three fluorescent reporters (Figure 9B). We hypothesize that this oxygen dependence is a result of active oxidative phosphorylation enabled by machinery in the crude cell lysate, which has been reported in previous studies.^23,39^ Specifically, inverted inner membrane vesicles formed upon homogenization are thought to carry functional components of the *E. coli* electron transport chain, which generate more ATP when oxygen is available as a terminal electron acceptor.^39^ Despite this drop in yield under anaerobic conditions, the high yields of *E. coli* lysate-based CFPS systems are such that our anaerobic CFPS still produces sufficient concentrations of protein to screen enzymatic activity in most cases.

### Synthesis of a functional, highly oxygen sensitive [4Fe-4S] cluster PFOR enzyme

As a proof of concept for the utility of an anaerobic CFPS platform, we used this system to produce an oxygen-sensitive [4Fe-4S] cluster protein—pyruvate:ferredoxin oxidoreductase (PFOR). PFOR plays an essential role in anaerobic metabolism, catalyzing the CoA-dependent oxidative decarboxylation of pyruvate into CO_2_ and acetyl-CoA, a key metabolic intermediate (Figure 10A).^40,41^ With the potential to tune energy fluxes, PFOR is a key target for drug development^42^ and metabolic engineering^43^ of anaerobes. However, PFORs remain difficult to study as they are highly oxygen sensitive, require iron-sulfur clusters for functional activity, and necessitate labor-intensive anaerobic protein purification to characterize protein function. To this end, we sought to demonstrate anaerobic CFPS of the PFOR from *Prevotella brevis* GA33 (henceforth referred to as PFOR33), an obligate anaerobic bacterium found in the rumen of domestic livestock facilitating the breakdown of plant biomass to propionate, acetate, and succinate (Figure S5).

**Figure 10.**
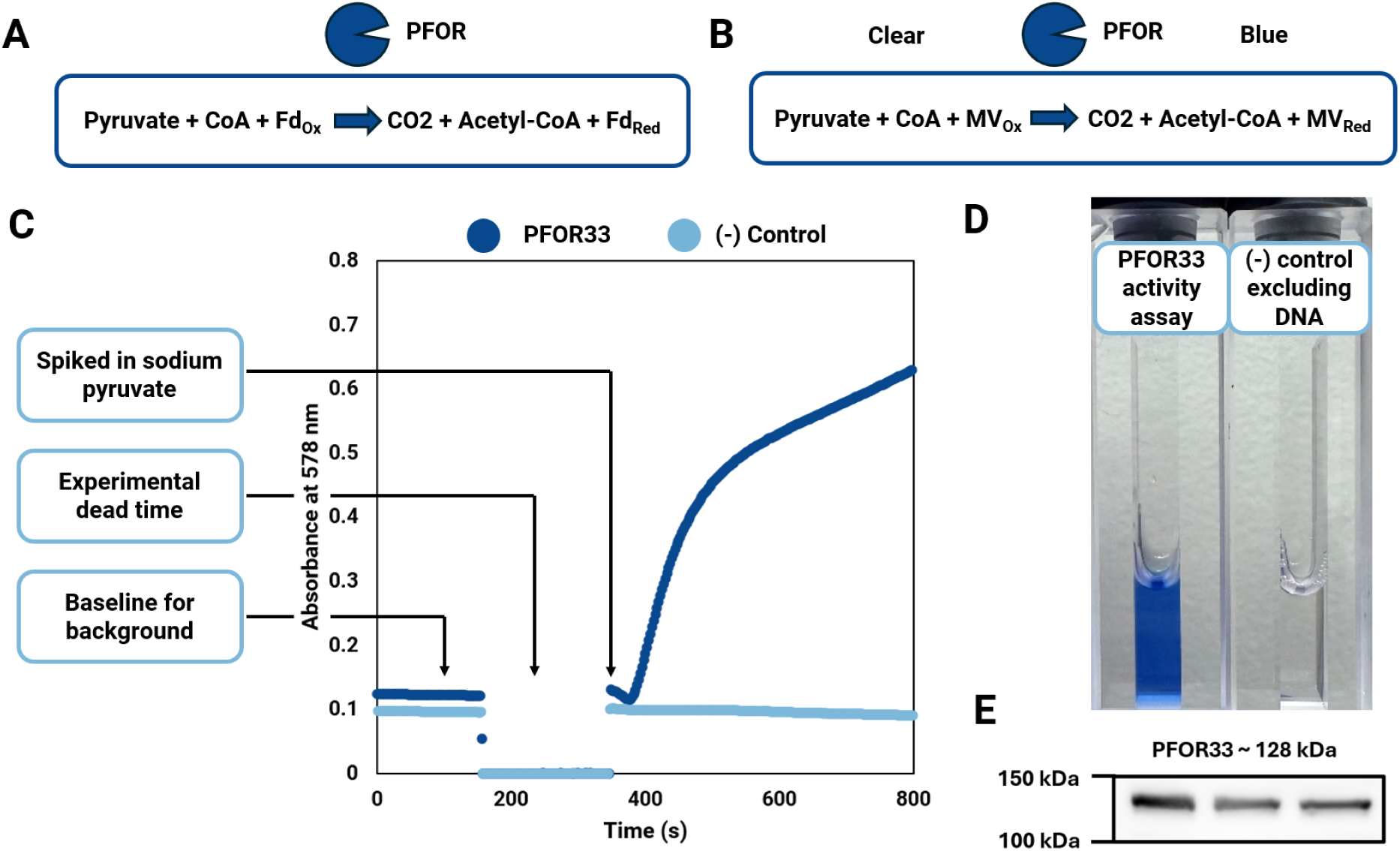
Demonstrating enzyme activity of pyruvate: ferredoxin oxidoreductase (PFOR), a [4Fe-4S] cluster enzyme produced via anaerobic cell-free protein synthesis. (A) PFOR reaction mechanism in which pyruvate is converted to acetyl-CoA and ferredoxin is reduced (Fd_ox_, oxidized ferredoxin; Fd_red_, reduced ferredoxin). (B) Reaction mechanism for a colorimetric PFOR activity assay in which methyl viologen acts as an artificial electron acceptor in place of ferredoxin (MV_ox_, oxidized methyl viologen, MV_red_, reduced methyl viologen). When methyl viologen is reduced in the reaction, it shifts in color from clear to blue, absorbing light at 578 nm. (C) Using a spectrophotometer we can monitor the change in absorbance at 578 nm and quantify the specific activity of PFOR33. The first region in this graph represents a baseline measurement of absorbance at this wavelength with all assay components excluding only sodium pyruvate. The second region represents experimental dead time in which the cuvette was removed from the spectrophotometer and sodium pyruvate was spiked in under constant flow of nitrogen gas. The last region represents the initiation of the reaction as PFOR33 catalyzed the oxidation of pyruvate and reduction of MV. We observed a specific activity of 5.2 ± 0.3 x 10^-4^ U/µL cell-free extract in which 1 U is defined as the oxidation of 1 μmol pyruvate and the reduction of 2 μmol methyl viologen, respectively per minute. (D) Visual representation of the color-shift of reduced methyl viologen between a positive and negative control. The negative control in this experiment contains all assay components including cell-free extract, however DNA encoding for PFOR33 was not supplied during anaerobic CFPS. (E) Cropped western blot showing the expected band for the N-terminal strep tag fused PFOR33 protein at a molecular weight of 128 kDa for three replicates.

We found that our anaerobic CFPS reactions produce functional PFOR33 with the supplementation of three additives (Table 2). Using our LyRA CFPS formulation as a base, we added three chemical compounds during rehydration, diluted with anoxic water. First, thiamine pyrophosphate was included because this is a requisite cofactor for PFOR activity. Second, to enable formation of the [4Fe-4S] clusters required for functional PFOR, we included ferrous ammonium sulfate and sodium sulfide as iron and sulfur sources, respectively. Successful production of PFOR33 was validated by western blot against the N-terminal Strep tag fused to the produced protein (Figure S6), with an estimated protein yield of 0.035 ± 0.004 µg/µL of cell-free extract (Figure S7). To confirm the functionality of the synthesized PFOR33, we used a methyl viologen assay.^44^ In this assay, the CoA-dependent oxidative decarboxylation of pyruvate by PFOR33 reduces methyl viologen (instead of ferredoxin) and initiates a color change from clear to blue, which can then be detected using absorbance (Figure 10B). We found that only LyRA CFPS reactions that include the DNA template for PFOR33 result in observable product formation, with a specific activity of 5.2 ± 0.3 x 10^-4^ U/µL cell-free extract (Figure 10C). Importantly, LyRA CFPS reactions that were supplemented with the same additives (thiamine pyrophosphate, ferrous ammonium sulfate, sodium sulfate), but no DNA template, did not yield any observable methyl viologen reduction.

**Table 2.** Chemical additives that allowed for [4Fe-4S] cluster assembly in vitro via anaerobic CFPS.

| Chemical | Concentration | Purpose |
|---|---|---|
| Ferrous ammonium sulfate (Fe(NH4)2(SO4)2) | 1 mM | Provides free Fe 2+ ions |
| Sodium sulfide (Na2S) | 0.8 mM | Provides free S 2- ions |
| Thiamine pyrophosphate (TPP) | 1 mM | Co-factor required for PFOR activity |

Together, these results demonstrate that minimal supplementation of our LyRA CFPS reactions allows production of a functional, oxygen-sensitive iron-sulfur cluster containing enzyme (Figure 10C-E).

Our LyRA method greatly simplified production of oxygen-sensitive [Fe-S] cluster enzymes via CFPS. In contrast to previous studies that rely on K12 strains of *E. coli*, minimal growth media, and partial or fully anaerobic growth for production of the lysate used in CFPS reactions, we used aerobically-grown BL21(DE3)* *E. coli* as the source of our cell lysate. This bypasses tedious media preparation steps and utilizes a strain optimized for protein production. Notably, LyRA still requires the use of a glovebox to set up CFPS reactions, whereas the PURE system supplemented with oxygen-scavenging enzymes developed by Po-Hsiang Wang *et al.*^17^ can synthesize [4Fe-4S] enzymes under aerobic conditions. However, the PURE anaerobic system requires the addition of flavin reductase, formate dehydrogenase, catalase, sodium formate, pyridoxal 5′-phosphate, NADH, FAD, and the mRNA encoding for their protein of interest just to produce the apo form of Fe-S containing enzymes. To initiate the maturation of holo-[Fe-S] cluster enzymes, FeSO_4_ and individually purified SUF subunits SufA, SufB, SufC, SufD, SufE, and SufS must be added. In comparison, our formulation only uses two chemical additives to produce a mature [4Fe-4S] cluster enzyme: ferrous ammonium sulfate and sodium sulfide. Moreover, we perform both transcription and translation in a single reaction rather than adding exogenous mRNA. Our LyRA workflow therefore builds on these bodies of work, streamlining the production of oxygen-sensitive enzymes in a one-pot reaction with few modifications from the widely used PANOx-SP CFPS formulation.

## Conclusions

Broader adoption of anaerobic microorganisms for use in industrial bioprocessing has been limited by a lack genetic tools to engineer mutant strains and the difficulty associated with characterizing oxygen-sensitive enzymes. In this study we demonstrate two easy-to-implement methods for anaerobic cell-free protein synthesis—a one-pot transcription and translation system to produce and screen the activity of oxygen-sensitive enzymes. Leveraging the widely used PANOx-SP formulation with lysate from BL21(DE3)* as our CFPS reaction base, we show that oxygen-free CFPS can be achieved by either sparging with nitrogen gas or lyophilizing and rehydrating CFPS reaction mixtures with anoxic water, a novel workflow we call LyRA. Because the CFPS formulation used here and the lyophilization of CFPS reactions are both widely adopted by many labs in the field, LyRA is broadly accessible to any lab already doing CFPS, and we expect this technology will be highly enabling to other research groups interested in anaerobic metabolism and cell-free expression. We illustrated this potential through the production of a functional [4Fe-4S] cluster pyruvate:ferredoxin oxidoreductase (PFOR) from the obligate anaerobic bacterium *Prevotella brevis* GA33,^22,45^ finding that only three reagents need to be added to the baseline formulation to produce a functional Fe-S cluster enzyme. Notably, this technology is highly modular, lending itself to high-throughput screens to characterize novel oxygen-sensitive proteins that may otherwise be inaccessible in their native cells. With our LyRA workflow, we can prepare large batches of anoxic CFPS reactions and store them for future use, retaining full activity four weeks after storage at -20 °C and -80 °C. This allows researchers to retrieve aliquots, rehydrate, add DNA, and produce highly oxygen sensitive proteins that can be characterized 24 hours after initiating the CFPS reaction.

## Acknowledgements

This material is based on work performed at the NSF BioFoundry for Extreme and Exceptional Fungi Archaea and Bacteria (ExFAB) and supported by the National Science Foundation under Award No. DBI-2400327. This work was partially supported by the U.S. Department of Energy, Office of Science and Technology (DE-SC0025492). This work was supported by the National Science Foundation Graduate Research Fellowship under Grant No. 2139319 to J.D.A and the University of California Santa Barbara Racial Justice Fellowship to J.D.A.

## Methods

### Molecular Cloning

#### Construction of pJL1-iRFP702 and pJL1-mFAP2a

The gene encoding for iRFP702 (GenBank: AGN32865) was codon-optimized for *E. coli* and 4-base pair overhangs for the Type IIS restriction enzyme, BsaI, were added to the 5’ and 3’ end of the gene sequence (SI Table 6). This gene block was synthesized by IDT, and codon-optimization was done using IDT’s Codon Optimization Tool. The gene encoding for mFAP2a was PCR-amplified from pET15b-TEV-mFAP2a^29^ (a gift from Tullman-Ercek lab) using primers oJDA001 and oJDA002 (SI Table 5). These primers added the requisite 4-base pair Golden Gate overhangs and BsaI restriction sites. The PCR product was then purified using a spin column (Promega Wizard™ SV Gel and PCR Cleanup System).

The pJL1-ggsfGFP plasmid (a gift from the Tullman-Ercek lab, sCEM100) was purified (E.Z.N.A.® Plasmid DNA Mini Kit I, (Q-spin, capless)) from *Escherichia coli* DH10B cells and used as the vector backbone for both constructs. In a 10 µL Golden Gate reaction, the pJL1-ggsfGFP plasmid and gene of interest were added at a 3:1 gene-of-interest: vector molar ratio to a total DNA content of 100-150 ng. The Type IIS restriction enzyme BsaI-HFv2 (NEB R3733), T4 ligase (NEB M0202T), and T4 Ligase Buffer were added to the reaction, which was then cycled 25 times between 37 °C for 2 minutes and 16 °C for 5 minutes, followed by a 5 minute incubation at 50 °C and a 5 minute incubation at 80 °C in a thermocycler. The Golden Gate reactions were then transformed into chemically competent DH10B cells and plated on LB-Agar plates supplemented with 50 µg/mL kanamycin. A successful Golden Gate reaction with this vector replaces the constitutively expressed sfGFP in pJL1-ggsfGFP with the new gene of interest; thus, after plating, white colonies are selected, as this indicates sfGFP was removed and replaced with the gene of interest. Selected colonies were used to inoculate cultures in LB-Miller supplemented with 50 µg/mL kanamycin, which were grown overnight and miniprepped. The two Golden Gate reactions yielded pJL1-iRFP702-Ecoli-Optim and pJL1-mFAP2a-Ecoli-Optim respectively, and were sequence-verified by whole plasmid sequencing at Plasmidsaurus Inc.

#### Construction of pET32a-Strep-PFOR33

The gene encoding pyruvate:ferredoxin oxidoreductase (PFOR33) of *Prevotella brevis* GA33 (T433DRAFT_00909, Locus Tag in IMG/MER) was amplified from genomic DNA and inserted into pASK-IBA7 (a gift from the Happe lab) at the site downstream of Strep-tag II, generating plasmid pASKIBA7-Strep-PFOR33. To purify genomic DNA, *P. brevis* GA33 (a gift from Hackmann lab) was grown in complex medium C at 37 °C.^22,44^ Medium C contains the following components per liter: 150 mL salt solution I, 150 mL salt solution II, 150 mL clarified rumen fluid, 2.5 g yeast extract, 10 g trypticase peptone, 6 g NaHCO_3_, 1 mL of 0.1% (w/v) resazurin solution, 1 g L-cysteine hydrochloride.^46^ Salt solution I contained 3 g/L K_2_HPO_4_ dissolved in distilled water. Salt solution II contained the following components per liter: 3 g KH_2_PO_4_, 6 g (NH_4_)_2_SO_4_, 6 g NaCl, 0.6 g MgSO_4_·7H_2_O, 0.6 g CaCl_2_·2H_2_O dissolved in water. Once the culture had reached stationary phase, 100 µL of *P. brevis* GA33 culture was pelleted, media was aspirated, the pellet was resuspended in 100 µL sterile water, and boiled for 10 min. This boiled cell mixture was then centrifuged, and the supernatant was used to amplify the gene using the Phusion PCR master Mix (NEB#M0531L). pASK-IBA7 backbone DNA was prepared by PCR with Phusion PCR Master Mix, digested with DpnI, and gel-purified. These DNA fragments were assembled using a Gibson Assembly kit (NEB #E2621L). The DNA fragment of PFOR33 with the N-terminal Strep-Tag II from pASKIBA7-Strep-PFOR33 was amplified by PCR and inserted into pET32a to replace the sequence between the start codon (ATG) and the stop codon (TAA) within the sequence downstream of the ribosome binding site but upstream of the T7 terminator, generating plasmid pET32a-Strep-PFOR33. The plasmids were sequence-verified via whole plasmid sequencing by Plasmidsaurus Inc.

### Cell Extract Preparation

Crude lysates from the *E. coli* strain BL21 (DE3)* were prepared in 1 L batches and grown in 2.5 L baffled Tunair^TM^ shake flasks (IBI Scientific^TM^ SS-5013).^21^ Cultures were inoculated from an overnight culture grown in LB-Miller to an optical density at 600 nm (OD600) of 0.08 and grown at 37 °C with shaking at 250 rpm in 2xYTPG media (10 g/L yeast extract, 16 g/L tryptone, 5 g/L sodium chloride, 7 g/L potassium phosphate dibasic, 3 g/L potassium phosphate monobasic, 18 g/L glucose, pH adjusted to 7.2 with potassium hydroxide). Once cultures reached OD600 between 0.6-0.8, expression of T7 RNA polymerase was induced by addition of 1 mM isopropyl β-D-1-thiogalactopyranoside (IPTG). The cultures then continued to grow at 37 °C with shaking until reaching an OD600 of 3, after which cells were harvested by centrifugation at 8,000xg for 10 minutes at 4 °C in an Avanti Centrifuge, rotor JLA 9.1000. Cell pellets were kept on ice for all subsequent steps. After removal of spent media, cell pellets were washed 3 times with S30 buffer (10 mM Tris acetate pH 8.2, 14 mM magnesium acetate, and 60 mM potassium acetate), where pellets were resuspended in 25 mL of S30 buffer per 0.5 L of culture. After each wash, cells were pelleted by centrifugation at 10,000xg for 2 minutes at 4 °C. Following wash steps, cell pellets were weighed, flash frozen in liquid nitrogen, and stored at -80 °C.

Prior to lysis, cells were thawed on ice and resuspended in 1 mL S30 buffer per gram of wet cell pellet. Cells were then lysed using a high-pressure homogenizer (Avestin EmulsiFlex-B15) with a single pass at a homogenizing pressure of 22,500 psi. Following lysis, cells were immediately cooled in a stainless-steel beaker on ice. Cell lysate then was aliquoted into 1.75 mL microcentrifuge tubes and centrifuged at 12,000xg for 10 minutes at 4 °C. Clarified lysate was aspirated from the top of each tube and transferred to new chilled 1.75 mL microcentrifuge tubes and centrifuged a second time at 12,000xg for 10 minutes at 4 °C. The resulting supernatant was then collected into a single stock, aliquoted into PCR tubes, flash frozen in liquid nitrogen, and stored at -80 °C.

### Cell-Free Protein Synthesis

All cell-free protein synthesis reactions in this work use the S30 PANOx-SP formulation as previously described elsewhere with final reagent concentrations and product numbers for each reagent noted in Tables S1 and S2.^21^ In brief, the base CFPS formulation includes the following reagents: 12 mM magnesium glutamate; 10 mM ammonium glutamate; 130 mM potassium glutamate; 1.2 mM ATP; 0.85 mM CTP; 0.85 mM UTP; 0.85 mM GTP; 0.03 mg/mL folonic acid; 0.17 mg/mL E. coli tRNA; 2 mM 20-amino acid mix; 33.33 mM PEP; 0.33 mM NAD; 0.27 mM CoA; 4 mM oxalic acid; 1 mM putrescine; 1.5 mM spermidine; 57 mM HEPES; 13.33 ng/µL plasmid DNA; 30% *E. coli* BL21(DE3)* cell extract by volume; and water.

Reactions were run at the 25 µL scale for data presented for degassing, lyophilization, and anaerobic vs aerobic proof of concept experiments. Reactions for testing the stability of lyophilized samples after storage were run at the 10 µL scale. Lastly, reactions used for the expression of PFOR33 were run at 30 µL scale to provide enough sample for activity assays and subsequent western blots; reagents required for Fe-S cluster reconstitution were added to PFOR33 CFPS reactions as described below (see “CFPS of PFOR33”).

All anaerobic reactions were conducted in a 384-well flat-bottom microplate and sealed with a gas impermeable seal (Microseal B Bio-Rad MSB1001B). Aerobic experiments were conducted in a 384-well flat-bottom microplate and sealed with a gas permeable seal that allows for the exchange of oxygen but prevents evaporation (Breathe-Easy® sealing membrane, Millipore Sigma Z380059). All cell-free reactions were incubated in sealed microplates at 30 °C for 24 hours without agitation.

### Preparation of anoxic water

In a 2 L Pyrex flask, 500 mL of Milli-Q purified double distilled water was boiled for 5-10 minutes in the microwave. The flask was then cooled on ice and bubbled with N_2_ gas for 30 minutes. Anoxic water was then transferred to 120 mL Wheaton serum bottles that were pre-flushed with nitrogen gas. Serum bottles were sealed with butyl rubber stoppers and crimped with a metal seal. Serum bottles containing anoxic water were then autoclaved for 20 minutes at 121 °C.

### Sparging cell-free extracts

To prepare sparged cell-free reactions, we began by preparing reagents aerobically. We then mixed cell-free reaction components under the constant flow of nitrogen gas. The flow rate of nitrogen gas and needle height were adjusted to allow for gentle bubbling to displace dissolved oxygen in the mixture. Careful adjustment of the flow rate is required, as higher flow rates cause the cell-free extract to foam over, resulting in a failed run (Figure S8). Reagents were added stepwise to a custom-blown glass cuvette whose opening seals with a rubber septum to create an air-tight vessel. In brief, we added the following reagents first: 15X salt solution, 15X NTP/tRNA Master mix, PEP, Amino acid mix, and 12.5X Reagent mix, notably excluding water and cell lysate. This mixture was bubbled for 5 minutes before S30 cell lysate was added. This mixture was bubbled for another 5 minutes. The glass cuvette was then sealed off with a rubber septum and set in the anaerobic chamber to rest for 4 hours, allowing bubbles to dissipate.

Once in the anaerobic chamber, anoxic water was added to the cell-free master mix to the final total reaction volume. The amount of water added was variable depending on the amount of water lost to evaporation during sparging. This master mix was then aliquoted into appropriate reaction volumes, and DNA and fluorogenic co-factors were added as necessary.

### Oxygen detection and scavenging strategies

#### Resorufin as an oxygen reporter

CFPS reactions were prepared by sparging with nitrogen gas as described above. A 1000X stock of resazurin (Sigma R7017-1G), at a concentration of 0.1% w/v or 4 mM, was prepared anaerobically and stored at 4 °C. The day of experimentation, a 10X working stock of resazurin was prepared anaerobically, diluted with anoxic water from the 1000X stock and added to the CFPS reaction at a final concentration of 4 µM in the anaerobic chamber. Resazurin functions as a redox indicator when reduced to resorufin. As such, resazurin was always used in combination with a reducing agent such as cysteine-HCl or DTT to enable its function as an oxygen indicator.

To assess the concentration of dissolved oxygen in our anaerobic CFPS reactions, we monitored resorufin fluorescence before and after exposing our samples to oxygen.

Specifically, we incubated CFPS samples under anaerobic conditions for 24 hours, allowing transcription and translation to proceed. We then measured fluorescence of resorufin with the following settings on the Tecan Spark microplate reader: excitation 559 nm, bandwidth 10 nm; emission 584 nm, bandwidth 10 nm. Resorufin fluorescence increases with the concentration of oxygen in solution. After taking an initial measurement of the sample under anoxic conditions, the seal was removed and samples were exposed to oxygen. Resorufin fluorescence was monitored as a function of time after exposure to oxygen over the course of 90 minutes, with one measurement taken per minute, shaking at a frequency of 180 rpm in the plate reader.

#### Cysteine-HCl and DTT as oxygen scavengers

CFPS reactions were prepared by sparging with nitrogen gas as described above. Stocks of cysteine-HCl (0.1 M, Fisher Scientific AC434850250) and DTT (0.1 M, Fisher Scientific BP172-5) were prepared anaerobically and stored at -20 °C. The day of experimentation, these were thawed and added to their respective CFPS reaction in the anaerobic chamber at a final concentration of 0.01 M.

### Lyophilization/rehydration/anaerobic (LyRA) CFPS

In this study, we screened 3 formulations to increase the stability of lyophilized cell-free reactions. The PANOx-SP BL21(DE3)* *E. coli* CFPS formulation is used as the base for all LyRA reactions as described in the “Cell-Free Protein Synthesis” section, with modifications described below.

#### Maltodextrin variation for LyRA (MD)

A stock of maltodextrin in water (300 mg/mL) was prepared fresh the day of lyophilization and added to CFPS reactions at a final concentration of 60 mg/mL as described previously.^33^

#### Maltodextrin-β-lactose variation for LyRA (MDBL)

Stocks of maltodextrin (300 mg/mL) and β-lactose (1 M) in water were prepared fresh the day of lyophilization and added to CFPS reactions to a final concentration of 60 mg/mL maltodextrin and 20 mM β-lactose, which are concentrations selected based on previous studies.^32,33^

#### Lyophilization and rehydration of CFPS samples with anoxic water

CFPS reactions were prepared aerobically on ice and aliquoted into PCR strip tubes or 1.5 mL microcentrifuge tubes. Samples and were then placed in a -80 °C freezer for 30 minutes or until frozen. All sample caps and lids were removed prior to lyophilization, and samples were then transferred to the Labconco FreeZone 2.5 Liter -50C Benchtop Freeze Dryer, with a 12-port drying chamber attachment. Lyophilization was performed overnight (16-20 hours) at 0.1 mbar, with the condenser set to -55 °C. After lyophilization, samples were visually inspected to ensure melting did not occur and samples were viable. Samples were then capped and stored in resealable zipper storage bags and placed at their appropriate storage temperature (room temperature, -20 °C, or -80 °C). Rehydration of lyophilized samples was done in the anaerobic chamber with a 95% N_2_/ 5% H_2_ headspace. Samples were rehydrated with anoxic water (prepared as described in “Preparation of anoxic water”), adding back 90% v/v water compared to the original volume. This assumes 10% of the CFPS reaction volume is solute that remains after lyophilization. This master mix was then aliquoted into appropriate reaction volumes where DNA and co-factors were added to individual reactions as appropriate.

### CFPS of PFOR33

Stocks of ferrous ammonium sulfate (0.1 M), sodium sulfide (0.08 M), and thiamine pyrophosphate (TPP) (0.05 M) were dissolved in anoxic water, flushed with nitrogen gas, and stored at -20 °C in 0.5 mL screw-top tubes. These reagents were added to lyophilized cell-free reactions after rehydration with anoxic water in the anaerobic chamber. Care was taken to mix by pipetting after the addition of each reagent to prevent the formation of insoluble products. Reagents were added to reach a final concentration of 1 mM ferrous ammonium sulfate, 0.8 mM sodium sulfide, and 1 mM TPP. Finally, samples were sealed and incubated at 30 °C for 24 hours to allow protein production to proceed.

### CFPS Yield Quantification of Fluorescent Reporters

#### Fluorescent measurements for iRFP702 and mFAP2a

iRFP702 ^27,28^ and mFAP2a ^29^ are oxygen-independent fluorescent reporter proteins used in this study to quantify relative CFPS yields under anaerobic conditions. Both reporters require co-factors (biliverdin and DFHBI for iRFP702 and mFAP2a, respectively) to bind to the reporter proteins to fluoresce. Both co-factors are light sensitive and were prepared in amber tubes. Stocks of biliverdin and DFHBI were prepared in DMSO at 25 mM and 10 mM respectively, and stored at -20 °C until use. In this study, biliverdin was added at a final concentration of 100 µM to all samples producing iRFP702, and DFHBI was added at a final concentration of 50 µM to all samples producing mFAP2a. To halt translation in these reactions after 24 hours, chloramphenicol was added at a final concentration of 100 µM. Fluorescence from iRFP702 was measured with the Tecan Spark microplate reader: excitation 673 nm, bandwidth 10 nm; emission 702 nm, bandwidth 10 nm. Fluorescence from mFAP2a was measured with the following settings: excitation 467 nm, bandwidth 10 nm; emission 505 nm, bandwidth 10 nm.

#### Comparing aerobic and anaerobic yields

Both aerobic and anaerobic CFPS reactions were prepared by sparging with nitrogen gas as described above. This large master mix was then split into two separate tubes for aerobic and anaerobic protein production. In the aerobic case, samples were moved to the benchtop and aliquoted into a 384-well flat-bottom microplate under atmospheric conditions, exposing all reactions to oxygen. DNA and co-factors were added before sealed with a gas permeable seal that allows for the exchange of oxygen but prevents evaporation (Breathe-Easy® sealing membrane, Millipore Sigma Z380059). Lastly, the samples were sprayed with pressurized air through the seal for 30 seconds to ensure reactions were fully oxygenated. In the anaerobic case, samples were moved into the anaerobic chamber to be aliquoted into a 384-well flat-bottom microplate, DNA and co-factors were added, and reactions were sealed with a gas impermeable seal (Microseal B Bio-Rad MSB1001B). After incubating both sets of samples for 24 hours at 30 °C, translation was halted with the addition of chloramphenicol at a final concentration of 100 µM. Fluorescence was measured for fluorescent reporter proteins after 90 minutes of exposure to oxygen, allowing the sfGFP chromophore to fully mature.

### Kinetic Tracking of sfGFP Chromophore Maturation

Anaerobic CFPS reactions were prepared by either sparging or lyophilization as described in sections “Sparging cell-free extracts,” and “Lyophilization/rehydration/anaerobic (LyRA) CFPS” respectively. After incubating samples for 24 hours at 30 °C in sealed 384-well plates, fluorescence measurements were taken using a Tecan Spark microplate reader. sfGFP fluorescence was measured using the following settings: excitation 485 nm, bandwidth 10 nm; emission 510 nm, bandwidth 10 nm. We then removed the seal and added chloramphenicol at a final concentration of 100 µM to halt translation. sfGFP maturation with oxygen exposure was then tracked by monitoring fluorescence for 90 minutes in the microplate reader shaking with the following settings: orbital, 3 mm amplitude, 180 rpm frequency.

### PFOR33 activity assay

CFPS reactions were prepared via LyRA as described in section “CFPS of PFOR33.” After allowing anaerobic protein synthesis to proceed for 24 hours, PFOR enzymatic activity was measured using a colorimetric assay in which methyl viologen (MV) acts as an electron acceptor in place of ferredoxin (Figure 10B). In this reaction, PFOR breaks down pyruvate and simultaneously reduces MV, triggering a colorimetric shift from clear to blue, which can be quantified by measuring absorbance at 578 nm (A578). The reaction mixture contained the following components, following the protocol outlined by Zhang *et al*. at a final volume of 500 µL: 50 mM Tris-Cl (pH 7.6), 10 mM MgCl_2_, 4 mM

Dithiothreitol (DTT), 0.2 mM Coenzyme A (CoA) lithium salt, 0.1 mM thiamine pyrophosphate (TPP), 2 mM methyl viologen (MV), 10 µL CFPS reaction, and 10 mM sodium pyruvate.^22^ The assay was performed at room temperature and prepared under constant flow of N_2_ at the benchtop to maintain anoxic conditions. All reaction components were mixed in a glass cuvette fitted to seal with a rubber stopper, excluding sodium pyruvate. Absorbance was monitored at 578 nm using a spectrophotometer to establish a baseline for background fluctuations in the assay mixture. Then sodium pyruvate was spiked in to initiate the reaction, and absorbance was measured at 578 nm for 15 minutes to monitor the reduction of MV.

### CFPS Yield Quantification of PFOR33

Semi-quantitative western blotting was used to approximate the yield of PFOR33 in crude CFPS reactions. Samples were prepared for western blotting as follows: 12.5 µL CFPS product or protein standard of known concentration; 25 µL SDS running buffer (3.03 g/L Tris base, 14.44 g/L glycine, 1 g/L SDS, pH 8.8); and 12.5 µL 4X Laemmli buffer containing β-mercaptoethanol (BME) as a reducing agent (10 g/L bromophenol blue, 80 g/L SDS, 0.2 M Tris, 20 vol% BME, 30 vol% glycerol). This reaction was mixed and incubated at 95 °C for 10 minutes, then cooled to room temperature. 10 µL of each sample was loaded in each experimental lane alongside commercial protein ladders (3 µL, Bio-Rad #1610374 and Bio-Rad #1610376) on a 10-lane 7.5 wt% polyacrylamide Tris-glycine gel. Protein bands were separated via electrophoresis by applying 60 V for 30 minutes then 200 V for 40 minutes. Samples were then transferred to Immun-Blot PVDF 0.2 µm membrane (Bio-Rad #1620177) by applying 70 V for 30 minutes in a wet-tank electrophoresis transfer system (Bio-Rad #1703930). Membranes were blocked for 1 hour at room temperature with 10 mL of 5 % w/v milk in tris-buffered saline (TBS, 20 mM Tris, 150 mM NaCl, pH 7.5) containing 0.05 vol% tween-20 (TBS-T). The primary anti-strep-tag II antibody (Abcam #Ab252885) at 1:5,000 dilution was diluted in 5 % w/v milk in TBS-T and incubated with the membrane overnight at 4 °C. The next day, the membrane was washed with 15 mL TBS-T for 2 minutes, repeated for a total of 4 washes. The membrane was then incubated with a horseradish peroxidase (HRP)-conjugated goat anti-Rat IgG secondary antibody (Abcam #Ab205720) at a 1:10,000 dilution, diluted in 5 % w/v milk in TBS-T, for 1 hour at room temperature. The membrane was then washed with 15 mL TBS-T for 2 minutes, repeated for a total of 4 washes, and a final rinse with Milli-Q water to increase the sensitivity of the detection reagent. Target protein bands were visualized with an enhanced chemiluminescence (ECL) detection kit (Bio-Rad #1705060S) and imaging was captured with the following settings on the Bio-Rad ChemiDoc: Chemiluminescence blot (647sp emission filter and no illumination light), Colorimetric blot (white epi-illumination using a 590/110 emissionfilter), and IR-Dye 680RD (far-red epi-illumination with a 715/30 emission filter). Image analysis and band densitometry were performed using ImageJ.

## Abbreviations

CFPS: Cell-free protein synthesis
sfGFP: Superfolder green fluorescent protein
iRFP702: Near IR fluorescent protein
mFAP2a: Mini fluorescence-activating protein 2a
LyRA: Lyophilization/rehydration/anaerobic workflow for anaerobic cell-free protein synthesis
MD: Maltodextrin
MDBL: Maltodextrin + β-lactose

## References

(1) Lu, Z.; Imlay, J. A. When Anaerobes Encounter Oxygen: Mechanisms of Oxygen Toxicity, Tolerance and Defence. Nat. Rev. Microbiol. 2021, 19 (12), 774–785. 10.1038/s41579-021-00583-y

(2) Podolsky, I. A.; Seppälä, S.; Lankiewicz, T. S.; Brown, J. L.; Swift, C. L.; O’Malley, M. A. Harnessing Nature’s Anaerobes for Biotechnology and Bioprocessing. Annu. Rev. Chem. Biomol. Eng. 2019, 10, 105–128. 10.1146/annurev-chembioeng-060718-030340

(3) Chen, W.; Mulchandani, A.; Deshusses, M. A. Environmental Biotechnology: Challenges and Opportunities for Chemical Engineers. AIChE J. 2005, 51 (3), 690–695. 10.1002/aic.10487

(4) Angenent, L. T.; Karim, K.; Al-Dahhan, M. H.; Wrenn, B. A.; Domíguez-Espinosa, R. Production of Bioenergy and Biochemicals from Industrial and Agricultural Wastewater. Trends Biotechnol. 2004, 22 (9), 477–485. 10.1016/j.tibtech.2004.07.001

(5) Amin, F. R.; Khalid, H.; El-Mashad, H. M.; Chen, C.; Liu, G.; Zhang, R. Functions of Bacteria and Archaea Participating in the Bioconversion of Organic Waste for Methane Production. Sci. Total Environ. 2021, 763, 143007. 10.1016/j.scitotenv.2020.143007

(6) Vees, C. A.; Neuendorf, C. S.; Pflügl, S. Towards Continuous Industrial Bioprocessing with Solventogenic and Acetogenic Clostridia: Challenges, Progress and Perspectives. J. Ind. Microbiol. Biotechnol. 2020, 47 (9–10), 753–787. 10.1007/s10295-020-02296-2

(7) Tan, Y.; Liang, J.; Lai, M.; Wan, S.; Luo, X.; Li, F. Advances in Synthetic Biology Toolboxes Paving the Way for Mechanistic Understanding and Strain Engineering of Gut Commensal *Bacteroides* Spp. and *Clostridium* Spp. Biotechnol. Adv. 2023, 69, 108272. 10.1016/j.biotechadv.2023.108272

(8) Imlay, J. A. Iron-Sulphur Clusters and the Problem with Oxygen. Mol. Microbiol. 2006, 59 (4), 1073–1082. 10.1111/j.1365-2958.2006.05028.x

(9) Yeh, Y.-H.; Sirk, S. J. Genetic Toolbox Development for Engineering *Bacteroides* and Other Bacterial Species. Curr. Opin. Microbiol. 2026, 90, 102709. 10.1016/j.mib.2026.102709

(10) Jewett, M. C.; Swartz, J. R. Mimicking the Escherichia Coli Cytoplasmic Environment Activates Long-Lived and Efficient Cell-Free Protein Synthesis. Biotechnol. Bioeng. 2004, 86 (1), 19–26. 10.1002/bit.20026

(11) Moore, S. J.; MacDonald, J. T.; Freemont, P. S. Cell-Free Synthetic Biology for in Vitro Prototype Engineering. Biochem. Soc. Trans. 2017, 45 (3), 785–791. 10.1042/BST20170011

(12) Jiang, L.; Zhao, J.; Lian, J.; Xu, Z. Cell-Free Protein Synthesis Enabled Rapid Prototyping for Metabolic Engineering and Synthetic Biology. Synth. Syst. Biotechnol. 2018, 3 (2), 90–96. 10.1016/j.synbio.2018.02.003

(13) Krüger, A.; Mueller, A. P.; Rybnicky, G. A.; Engle, N. L.; Yang, Z. K.; Tschaplinski, T. J.; Simpson, S. D.; Köpke, M.; Jewett, M. C. Development of a Clostridia-Based Cell-Free System for Prototyping Genetic Parts and Metabolic Pathways. Metab. Eng. 2020, 62, 95–105. 10.1016/j.ymben.2020.06.004

(14) Sun, Z. Z.; Yeung, E.; Hayes, C. A.; Noireaux, V.; Murray, R. M. Linear DNA for Rapid Prototyping of Synthetic Biological Circuits in an Escherichia Coli Based TX-TL Cell-Free System. ACS Synth. Biol. 2014, 3 (6), 387–397. 10.1021/sb400131a

(15) Boyer, M. E.; Wang, C.-W.; Swartz, J. R. Simultaneous Expression and Maturation of the Iron-Sulfur Protein Ferredoxin in a Cell-Free System. Biotechnol. Bioeng. 2006, 94 (1), 128–138. 10.1002/bit.20830

(16) Boyer, M. E.; Stapleton, J. A.; Kuchenreuther, J. M.; Wang, C.; Swartz, J. R. Cell-free synthesis and maturation of [FeFe] hydrogenases. Biotechnol. Bioeng. 2008, 99 (1), 59–67. 10.1002/bit.21511

(17) Wang, P.-H.; Nishikawa, S.; McGlynn, S. E.; Fujishima, K. One-Pot De Novo Synthesis of [4Fe-4S] Proteins Using a Recombinant SUF System under Aerobic Conditions. ACS Synth. Biol. 2023, 12 (10), 2887–2896. 10.1021/acssynbio.3c00155

(18) Tamiev, B. D.; Dopp, J. L.; Reuel, N. F. Anaerobic Conditioning of E. Coli Cell Lysate for Enhanced In Vitro Protein Synthesis. ACS Synth. Biol. 2021, 10 (4), 716–723. 10.1021/acssynbio.0c00501

(19) Lin, X.; Zhou, C.; Zhu, S.; Deng, H.; Zhang, J.; Lu, Y. O2-Tuned Protein Synthesis Machinery in Escherichia Coli-Based Cell-Free System. Front. Bioeng. Biotechnol. 2020, 8. 10.3389/fbioe.2020.00312

(20) Dopp, B. J. L.; Tamiev, D. D.; Reuel, N. F. Cell-Free Supplement Mixtures: Elucidating the History and Biochemical Utility of Additives Used to Support *in Vitro* Protein Synthesis in *E. Coli* Extract. Biotechnol. Adv. 2019, 37 (1), 246–258. 10.1016/j.biotechadv.2018.12.006

(21) Silverman, A. D.; Kelley-Loughnane, N.; Lucks, J. B.; Jewett, M. C. Deconstructing Cell-Free Extract Preparation for in Vitro Activation of Transcriptional Genetic Circuitry. ACS Synth. Biol. 2019, 8 (2), 403–414. 10.1021/acssynbio.8b00430

(22) Zhang, B.; Lingga, C.; De Groot, H.; Hackmann, T. J. The Oxidoreductase Activity of Rnf Balances Redox Cofactors during Fermentation of Glucose to Propionate in Prevotella. Sci. Rep. 2023, 13 (1), 16429. 10.1038/s41598-023-43282-9

(23) Olsen, M. L.; Copeland, C. E.; Sundberg, C. A.; Aw, R.; Shaver, Z. M.; Rao, G.; Swartz, J. R.; Karim, A. S.; Jewett, M. C. Design-Driven Optimization of Low-Cost Reagent Formulations for Reproducible and High-Yielding Cell-Free Gene Expression. Nat. Commun. 2026, 17 (1), 3478. 10.1038/s41467-026-69605-8

(24) Pédelacq, J.-D.; Cabantous, S.; Tran, T.; Terwilliger, T. C.; Waldo, G. S. Engineering and Characterization of a Superfolder Green Fluorescent Protein. Nat. Biotechnol. 2006, 24 (1), 79–88. 10.1038/nbt1172

(25) Iizuka, R.; Yamagishi-Shirasaki, M.; Funatsu, T. Kinetic Study of de Novo Chromophore Maturation of Fluorescent Proteins. Anal. Biochem. 2011, 414 (2), 173–178. 10.1016/j.ab.2011.03.036

(26) Craggs, T. D. Green Fluorescent Protein: Structure, Folding and Chromophore Maturation. Chem. Soc. Rev. 2009, 38 (10), 2865–2875. 10.1039/B903641P

(27) Shcherbakova, D. M.; Verkhusha, V. V. Near-Infrared Fluorescent Proteins for Multicolor in Vivo Imaging. Nat. Methods 2013, 10 (8), 751–754. 10.1038/nmeth.2521

(28) Hooker, C. A.; Hanafy, R.; Hillman, E. T.; Muñoz Briones, J.; Solomon, K. V. A Genetic Engineering Toolbox for the Lignocellulolytic Anaerobic Gut Fungus Neocallimastix Frontalis. ACS Synth. Biol. 2023, 12 (4), 1034–1045. 10.1021/acssynbio.2c00502

(29) Klima, J. C.; Doyle, L. A.; Lee, J. D.; Rappleye, M.; Gagnon, L. A.; Lee, M. Y.; Barros, E. P.; Vorobieva, A. A.; Dou, J.; Bremner, S.; Quon, J. S.; Chow, C. M.; Carter, L.; Mack, D. L.; Amaro, R. E.; Vaughan, J. C.; Berndt, A.; Stoddard, B. L.; Baker, D. Incorporation of Sensing Modalities into de Novo Designed Fluorescence-Activating Proteins. Nat. Commun. 2021, 12 (1), 856. 10.1038/s41467-020-18911-w

30. (30) Spectrum [Resorufin] | AAT Bioquest. https://www.aatbio.com/fluorescence-excitation-emission-spectrum-graph-viewer/resorufin (accessed 2026-11-05)

(31) Smith, M. T.; Berkheimer, S. D.; Werner, C. J.; Bundy, B. C. Lyophilized Escherichia Coli-Based Cell-Free Systems for Robust, High-Density, Long-Term Storage. BioTechniques 2014, 56 (4), 186–193. 10.2144/000114158

(32) Shivakumar, T.; Clark, J.; Goode, A.; Anyanwu, V. E.; Williams, P. M. A Design of Experiments Approach for Enhancing Room Temperature Stability of a Lyophilised and Paper-Based Bacterial Cell-Free System. Bioengineering 2025, 12 (3), 223. 10.3390/bioengineering12030223

(33) Warfel, K. F.; Williams, A.; Wong, D. A.; Sobol, S. E.; Desai, P.; Li, J.; Chang, Y.-F.; DeLisa, M. P.; Karim, A. S.; Jewett, M. C. A Low-Cost, Thermostable, Cell-Free Protein Synthesis Platform for On-Demand Production of Conjugate Vaccines. ACS Synth. Biol. 2022, 12 (1), 95–107. 10.1021/acssynbio.2c00392

(34) Jung, J. K.; Rasor, B. J.; Rybnicky, G. A.; Silverman, A. D.; Standeven, J.; Kuhn, R.; Granito, T.; Ekas, H. M.; Wang, B. M.; Karim, A. S.; Lucks, J. B.; Jewett, M. C. At-Home, Cell-Free Synthetic Biology Education Modules for Transcriptional Regulation and Environmental Water Quality Monitoring. ACS Synth. Biol. 2023, 12 (10), 2909–2921. 10.1021/acssynbio.3c00223

(35) Paiva, Y. F.; de Figueirêdo, R. M. F.; Queiroz, A. J. de M.; dos Santos, F. S.; Amadeu, L. T. S.; de Lima, A. G. B.; de Lima, T. L. B.; da Silva, W. P.; Moura, H. V.; Silva, E. T. de V.; Costa, C. C.; Galdino, P. O.; Gomes, J. P.; Leão, D. A. S. Maltodextrin as a Drying Adjuvant in the Lyophilization of Tropical Red Fruit Blend. Molecules 2023, 28 (18), 6596. 10.3390/molecules28186596

(36) Alibekov, R. S.; Aitbayeva, A. Z.; Khamitova, B. M.; Azimov, A. M.; Sulaiman, A.; Mustapa Kamal, S. M.; Taip, F. S. Effects of Maltodextrin in Freeze Drying on the Physical and Functional Properties of Different Type of Milk Powder. Cogent Food Agric. 2025, 11 (1), 2473540. 10.1080/23311932.2025.2473540

(37) Corveleyn, S.; Remon, J. P. Maltodextrins as Lyoprotectants in the Lyophilization of a Model Protein, LDH. Pharm. Res. 1996, 13 (1), 146–150. 10.1023/a:1016006106821

38. Calhoun, K. A.; Swartz, J. R. Energy Systems for ATP Regeneration in Cell-Free Protein Synthesis Reactions. In In Vitro Transcription and Translation Protocols; Grandi, G., Ed.; Humana Press: Totowa, NJ, 2007; pp 3–17. 10.1007/978-1-59745-388-2_1

(39) Jewett, M. C.; Calhoun, K. A.; Voloshin, A.; Wuu, J. J.; Swartz, J. R. An Integrated Cell-Free Metabolic Platform for Protein Production and Synthetic Biology. Mol. Syst. Biol. 2008, 4, 220. 10.1038/msb.2008.57

(40) Ferry, J. G. Acetate Metabolism in Anaerobes from the Domain Archaea. Life 2015, 5 (2), 1454–1471. 10.3390/life5021454

(41) Krivoruchko, A.; Zhang, Y.; Siewers, V.; Chen, Y.; Nielsen, J. Microbial Acetyl-CoA Metabolism and Metabolic Engineering. Metab. Eng. 2015, 28, 28–42. 10.1016/j.ymben.2014.11.009

(42) Samuelson, J. Why Metronidazole Is Active against Both Bacteria and Parasites. Antimicrob. Agents Chemother. 1999, 43 (7), 1533–1541. 10.1128/aac.43.7.1533

(43) Ku, J. T.; Chen, A. Y.; Lan, E. I. Metabolic Engineering Design Strategies for Increasing Acetyl-CoA Flux. Metabolites 2020, 10 (4), 166. 10.3390/metabo10040166

(44) Zhang, B.; Hrdy, I.; Tachezy, J.; Gao, Y.; Williams, S. M.; Fulcher, J. M.; Munoz, N.; Burnet, M.; Baker, S. E.; O’Malley, M. A. The Anaerobic Fungus Caecomyces Churrovis Produces H2 via a Non-Bifurcating NADH-Dependent Enzyme Complex. mBio 2026, 17 (9), e01643–26. 10.1128/mbio.01643-26

(45) Avgustin, G.; Wallace, R. J.; Flint, H. J. Phenotypic Diversity among Ruminai Isolates of Prevotella Ruminicola: Proposal of Prevotella Brevis Sp. Nov., Prevotella Bryantii Sp. Nov., and Prevotella Albensis Sp. Nov. and Redefinition of Prevotella Ruminicola. Int. J. Syst. Evol. Microbiol. 1997, 47 (2), 284–288. 10.1099/00207713-47-2-284

(46) Davies, D. R.; Theodorou, M. K.; Lawrence, M. I. G.; Trinci, A. P. J. Distribution of Anaerobic Fungi in the Digestive Tract of Cattle and Their Survival in Faeces. Microbiology 1993, 139 (6), 1395–1400. 10.1099/00221287-139-6-1395

