## Supplemental Information for "An anaerobic cell-free protein synthesis formulation for the *in vitro* study of oxygen sensitive proteins"

### Supporting Information:

**Supporting Figures S1-S8**

**Supporting Tables S1-S6**

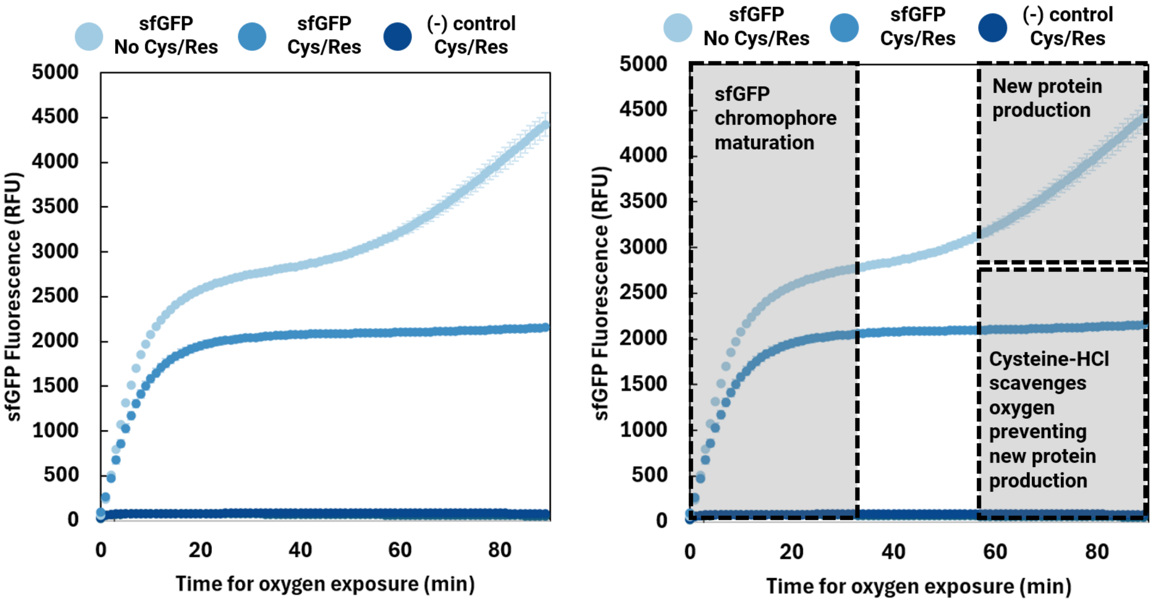


***Figure S1.*** *sfGFP chromophore maturation after exposure to oxygen, protein synthesis continued as oxygen diffuses into the reaction. Chloramphenicol was not added to halt these reactions after oxygen exposure, as a result, some reactions (those not containing Cysteine-HCl) continued to produce protein after 1 hour. Error bars represent the standard error of the mean (n=4).*

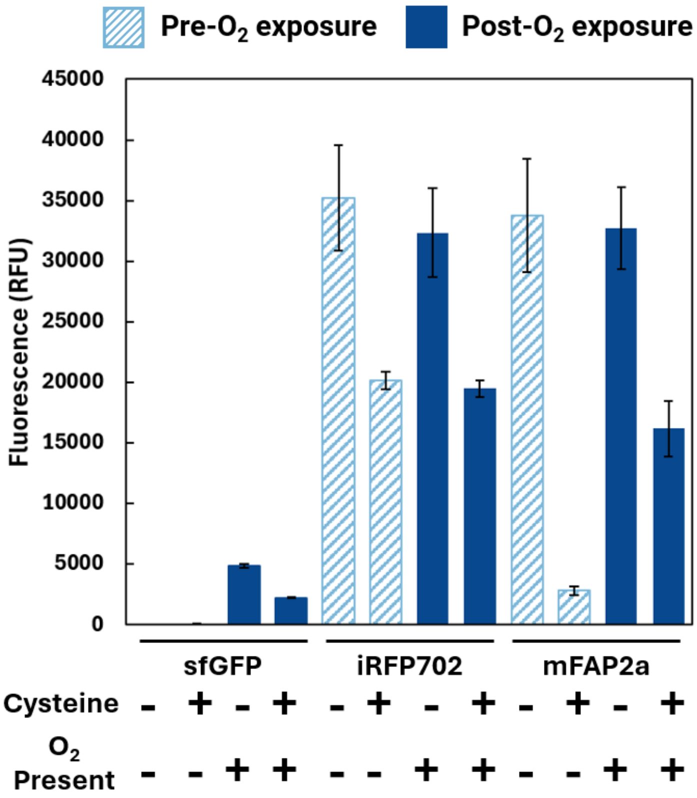


***Figure S2.*** *Effect of cysteine-HCl on protein production in anaerobic CFPS. The relative fluorescence of 3 fluorescent reporter proteins (sfGFP, iRFP702, and mFAP2a) above background is shown as a relative quantification for protein produced with and without cysteine-HCl, a reducing agent and oxygen scavenger. Cysteine-HCl consistently lowered CFPS yields across all three fluorescent reporter proteins. After incubating for 24 hours at 30 °C in sealed 384-well plates, fluorescence measurements were taken. Then, the seal was removed, translation was halted with the addition of 100 µM chloramphenicol, and fluorescence was measured again after 90 minutes of exposure to oxygen. This data represents fluorescence before and after oxygen exposure with sfGFP fluorescence maturing after exposure and iRFP702 and mFAP2a fluorescence remaining constant. Error bars represent ±1 standard error of the mean for n=4 sample size.*

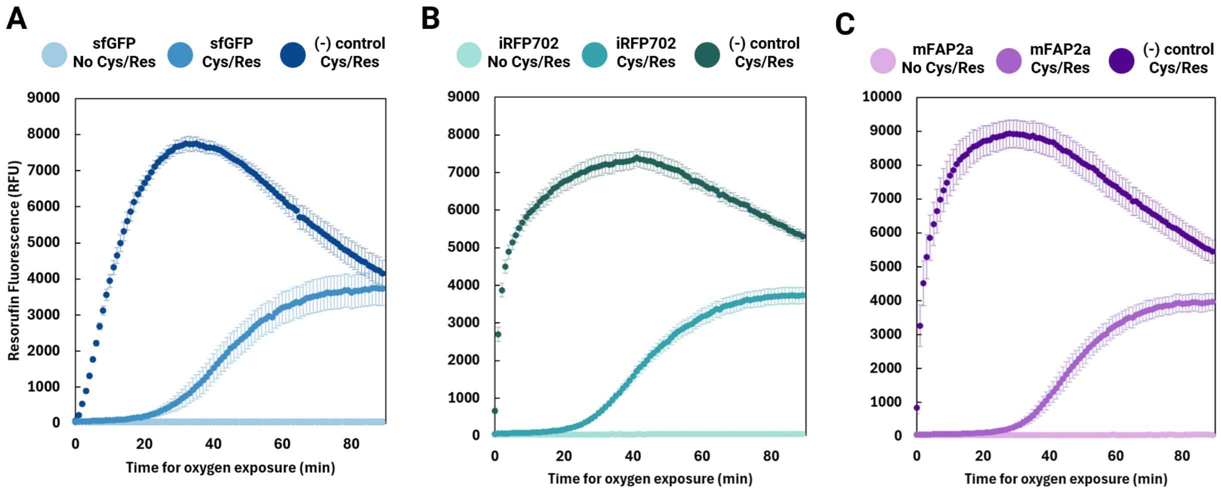


***Figure S3***. *Resorufin serves as an oxygen indicator in anaerobic cell-free reactions. For all fluorescent reporter proteins tested, we observed the same trend for resorufin fluorescence increasing as anaerobic CFPS reactions are exposed to oxygen. Resorufin fluorescence in response to oxygen exposure lags when protein is produced in the reaction compared to negative controls (that do not contain DNA).*

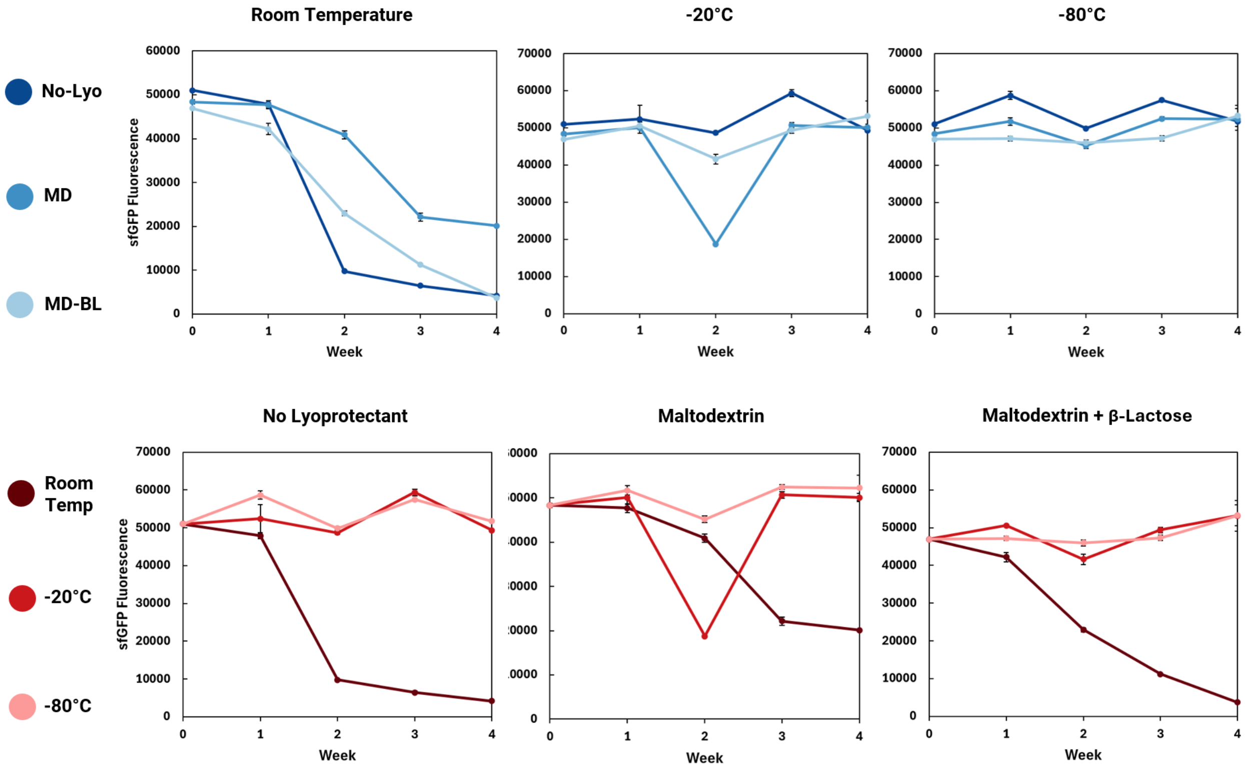


***Figure S4.*** *Optimization of formulation and storage conditions for aerobic lyophilized CFPS reactions. CFPS reactions were lyophilized in a large batch of aliquots and stored at room temperature, -20 °C, or -80 °C for up to 4 weeks. Each week, an aliquot was removed and rehydrated to determine the level of protein synthesis activity retained after long-term storage at the indicated temperature.*

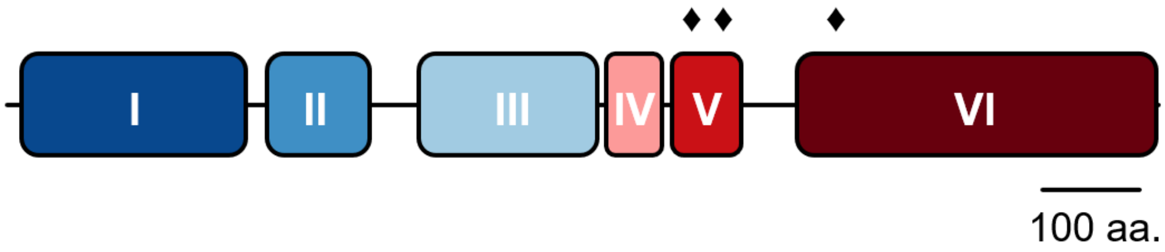


***Figure S5.*** *Diagram of predicted domains in PFOR33 from Prevotella brevis GA33. Domain I: thiamin diphosphate-binding fold; Domain II: pyruvate:ferredoxin oxidoreductase core domain II; Domain III: PFOR catalytic domain; Domain IV: EKR domain; Domain V: 4Fe-4S dicluster domain; Domain VI: C-terminal thiamine pyrophosphate binding domain. ♦: [4Fe-4S] cluster. The cysteines in [4Fe-4S] cluster 1 are at residues 692, 695, 698, and 759. Cysteines in [4Fe-4S] cluster 2 are at residues 702, 749, 752, and 755. Cysteines in [4Fe-4S] cluster 3 are at residues 824, 827, 852, and 1087. PFOR33 is 1192 amino acids long.*

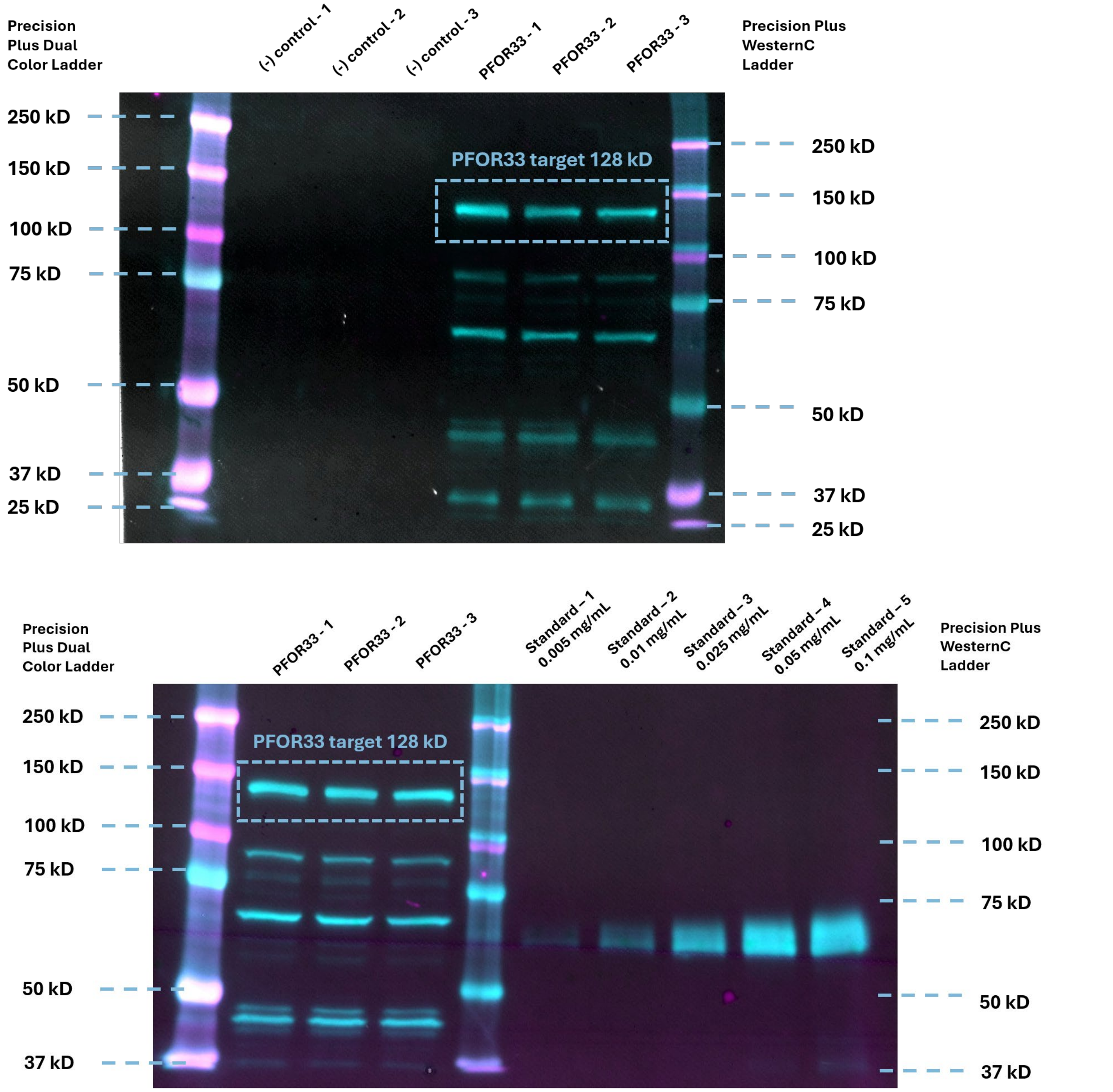


***Figure S6.*** *Uncropped anti-strep western blots confirming the presence of N-terminal strep tag fused PFOR33. In the top panel from left to right, (1) Precision Plus Dual Color Ladder, (2-4) negative controls for our CFPS reactions omitting DNA in triplicate, (5-7) anaerobic CFPS reactions that produced PFOR33 in triplicate, (8) Precision Plus WesternC Ladder. This confirmed the presence of our target protein ~128 kDa only in experimental samples where DNA encoding for PFOR33 was added. In the bottom panel from left to right, (1) Precision Plus Dual Color Ladder, (2-4) anaerobic CFPS reactions that produced PFOR33 in triplicate, (5) Precision Plus WesternC Ladder, (6-10) ladder of a purified strep-tagged protein ~60 kDa at concentrations between 0.005*

*mg/mL and 0.1 mg/mL. This was used to relatively quantify the yield of our anaerobic CFPS reaction at producing PFOR33.*

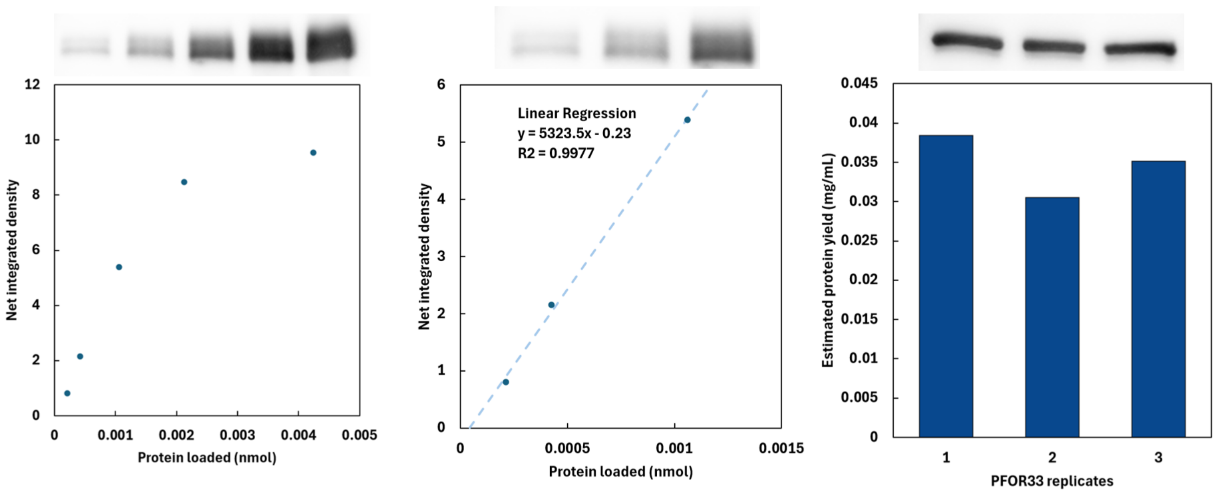


**Figure S7.** Quantification of PFOR33 yield from LyRA reactions. Using the western blots in Figure S6, we quantified the relationship between band density and the concentration of protein loaded in each lane. We then estimated the yield of PFOR33 to be 0.035 ± 0.004 µg PFOR33/µL of cell-free extract.

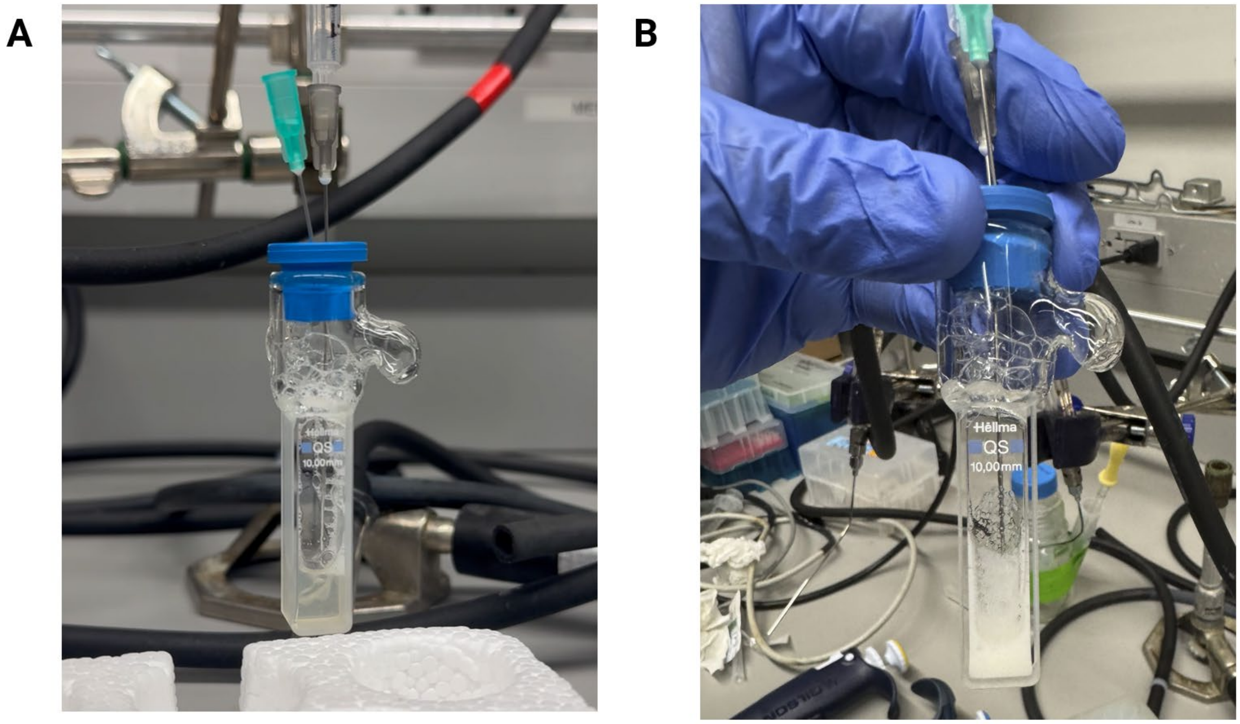


**Figure S8.** Visual comparison of a successful (A) versus unsuccessful (B) sparging experiment in which too much unwanted foam negatively impacts protein yield in CFPS reactions.

**Table S1. Components of the base CFPS reaction formulation used in this study.** The final component concentrations are given in mM unless otherwise stated.

| Component | Concentration | Vendor | Catalog Number |
|---|---|---|---|
| Magnesium glutamate | 12 | Millipore Sigma | 49605-250G |
| Ammonium Glutamate | 10 | Biosynth | FG28929 |
| Potassium glutamate | 130 | Millipore Sigma | G1501-100G |
| ATP | 1.2 | Millipore Sigma | A2383-5G |
| CTP | 0.85 | Millipore Sigma | C1506-1G |
| UTP | 0.85 | Millipore Sigma | U6750-500MG |
| GTP | 0.85 | Millipore Sigma | G8877-1G |
| Folinic acid | 0.03 mg/mL | Millipore Sigma | 47612-250MG |
| tRNA | 0.17 mg/mL | Millipore Sigma | R1753-500UN |
| 20 amino acid mix | 2 | See Table S2 | See Table S2 |
| PEP | 33.33 | Millipore Sigma | 860077-1G |
| NAD | 0.33 | Millipore Sigma | N8535-15VL |
| CoA | 0.27 | Millipore Sigma | C3144-500MG |
| Oxalic acid | 4 | Millipore Sigma | P0963-100G |
| Putrescine | 1 | Millipore Sigma | P7505-25G |
| Spermidine | 1.5 | Millipore Sigma | S2626-5G |
| HEPES | 57 | Millipore Sigma | H3375-1KG |
| Plasmid DNA | 13.33 ng/uL | | |
| E. coli cell extract | 30 % v/v | | |

**Table S2. Product numbers for the 20 amino acid mix used in the CFPS reaction formulation from this study.** All amino acids were added at a final concentration of 50 mM. This 50mM amino acid stock mix was stored at -80 °C until use and diluted to a final concentration of 2 mM in each CFPS reaction.

| Component | Vendor | Catalog Number |
|---|---|---|
| L-Valine | Millipore Sigma | V0500-25G |
| L-Tryptophan | Millipore Sigma | T0254-5G |
| L-Phenylalanine | Millipore Sigma | P2126-100G |
| L-Isoleucine | Millipore Sigma | I2752-10G |
| L-Leucine | Millipore Sigma | L8000-25G |
| L-Cysteine | Millipore Sigma | C7352-25G |
| L-Methionine | Millipore Sigma | M9625-25G |
| L-Alanine | Millipore Sigma | A7627-1G |
| L-Arginine | Millipore Sigma | A8094-25G |
| L-Asparagine | Millipore Sigma | A0884-25G |
| L-Aspartic acid | Millipore Sigma | A7219-100G |
| L-Glutamic acid, K salt | Millipore Sigma | G1501-100G |
| L-Glycine | Millipore Sigma | G7126-100G |
| L-Glutamine | Millipore Sigma | G3126-100G |
| L-Histidine | Millipore Sigma | H8000-10G |
| L-Lysine | Millipore Sigma | L5501-1G |
| L-Proline | Millipore Sigma | P0380-100G |
| L-Serine | Millipore Sigma | S4500-1G |
| L-Threonine | Millipore Sigma | T8625-10G |
| L-Tyrosine | Millipore Sigma | T8566-25G |

**Table S3.** Description of strains and plasmids used in this study

| Strain or Plasmid | Description | Study/Supplier |
|---|---|---|
| BL21 Star (DE3) | *E. coli* B strain optimized for T7-promoter-based protein expression systems | ThermoFisher Scientific(C601003) |
| DH10B | *E. coli* strain used for cloning DNA constructs | ThermoFisher Scientific (EC0113) |
| sCEM099: pJL1-sfGFP | sfGFP expressed under a T7 promoter in the pJL1 expression vector | Stark et al, Science Advances, 2018 (Addgene 102634) |
| sCEM100: pJL1-ggsfGFP | pJL1 Golden Gate entry vector, where constitutively expressed sfGFP is surrounded by BsaI sites and removed/replaced upon successful assembly | |
| sCEM241: pJL1-iRFP702-Ecoli-Optim | Oxygen-independent fluorescent reporter, iRFP702, codon optimized for *E. coli*, in the pJL1 expression vector | This study |
| sCEM242: pJL1-mFAP2a-Ecoli-Optim | Oxygen-independent fluorescent reporter, mFAP2a, codon optimized for *E. coli*, in the pJL1 expression vector | This study |
| sCEM562: pET32a-Strep-PFOR33 | Pyruvate:ferredoxin oxidoreductase (PFOR33) of *Prevotella brevis* GA33 with a N-terminal Strep-Tag II in the pET32a expression vector | This study |

**Table S4.** Plasmids used in this study

| Name | Plasmid | Origin | Resistance | Plasmid Assembly Method |
|---|---|---|---|---|
| sCEM099 | pJL1-sfGFP | colE1 | Kanamycin | Gift from Tullman-Ercek lab |
| sCEM100 | pJL1-ggsfGFP | colE1 | Kanamycin | Gift from Tullman-Ercek lab |
| sCEM241 | pJL1-iRFP702-Ecoli-Optim | colE1 | Kanamycin | Golden gate |
| sCEM242 | pJL1-mFAP2a-Ecoli-Optim | colE1 | Kanamycin | Golden gate |
| sCEM562 | pET32a-Strep-PFOR33 | colE1 | Ampicillin/Carbenicillin | Gibson assembly |

**Table S5.** Primers used in this study

| Name | Purpose | Description | Sequence (5' to 3') |
|---|---|---|---|
| oJDA001 | Golden gate assembly of pJL1-mFAP2a-Ecoli-Optim | Amplify mFAP2a with BsaI overhang | attaGGTCTCaCATGTCACGCGCCG |
| oJDA002 | Golden gate assembly of pJL1-mFAP2a-Ecoli-Optim | Amplify mFAP2a with BsaI overhang | taatGGTCTCaTTTATTCCTGACGCTGCA |
| oBZ046 | PCR amplification | Amplify PFOR33 from genome of *P. brevis* GA33 | ATGGCTAAAGAAAAGAAGTTTATCACC |
| oBZ047 | PCR amplification | Amplify PFOR33 from genome of *P. brevis* GA33 | TTACTCAGTCCAATCTTCCTGAGA |
| oBZ048 | Gibson Assembly of pASKIBA7 | Add overlap to PFOR33 for pASKIBA7 | CCCGCAGTTCGAAAAAGCTAAAGA AAAGAAGTTTATCACCTGTGATGG |
| oBZ049 | Gibson Assembly of pASKIBA7 | Add overlap to PFOR33 for pASKIBA7 | GGTCAAGCTTAGTTAGATATCATTAC TCAGTCCAATCTTCCTGAGACT |
| oBZ011 | Gibson Assembly of pASKIBA7 | Amplify backbone of pASKIBA7 | TGATATCTAACTAAGCTTGACCTGTG AAGTG |
| oBZ012 | Gibson Assembly of pASKIBA7 | Amplify backbone of pASKIBA7 | TTTTTCGAACTGCGGGTGGC |
| oBZ152 | Gibson Assembly of pET32a-Strep-PFOR33 | Amplify Strep-PFOR33 and add overlap to pET32a | TAGAAATAATTTTGTTTAACTTTAAGAAGG AGATATACATATGGCTAGCTGGAGCCAC |
| oBZ153 | Gibson Assembly of pET32a- | Amplify Strep-PFOR33 and | AACTCAGCTTCCTTTCGGGCTTTG TTACTCAGTCCAATCTTCCTGAGACTTGC |

| | | | |
|---|---|---|---|
| | Strep-PFOR33 | add overlap to pET32a | |
| oBZ029 | Gibson Assembly of pET32a-Strep-PFOR33 | Amplify backbone of pET32a | CAAAGCCCGAAAGGAAGCTGAG |
| oBZ154 | Gibson Assembly of pET32a-Strep-PFOR33 | Amplify backbone of pET32a | ATGTATATCTCCTTCTTAAAGTTAAACAAAATTATTTCTAGAGGGG |

**Table S6.** Gene sequences for proteins used in this study

| Gene | Sequence (5' to 3') |
|---|---|
| sfGFP | ATGAGCAAAGGTGAAGAACTGTTTACCGGCGTTGTGCCGATTCTG<br>GTGGAACTGGATGGCGATGTGAACGGTCACAAATTCAGCGTGCGT<br>GGTGAAGGTGAAGGCGATGCCACGATTGGCAAACTGACGCTGAAA<br>TTTATCTGCACCACCGGCAAACTGCCGGTGCCGTGGCCGACGCTG<br>GTGACCACCCTGACCTATGGCGTTCAGTGTTTTAGTCGCTATCCGG<br>ATCACATGAAACGTCACGATTTCTTTAAATCTGCAATGCCGGAAGG<br>CTATGTGCAGGAACGTACGATTAGCTTTAAAGATGATGGCAAATATA<br>AAACGCGCGCCGTTGTGAAATTTGAAGGCGATACCCTGGTGAACC<br>GCATTGAACTGAAAGGCACGGATTTTAAAGAAGATGGCAATATCCT<br>GGGCCATAAACTGGAATACAACTTTAATAGCCATAATGTTTATATTAC<br>GGCGGATAAACAGAAAAATGGCATCAAAGCGAATTTTACCGTTCGC<br>CATAACGTTGAAGATGGCAGTGTGCAGCTGGCAGATCATTATCAGC<br>AGAATACCCCGATTGGTGATGGTCCGGTGCTGCTGCCGGATAATCA<br>TTATCTGAGCACGCAGACCGTTCTGTCTAAAGATCCGAACGAAAAA<br>GGCACGCGGGACCACATGGTTCTGCACGAATATGTGAATGCGGCA<br>GGTATTACGTGGAGCCATCCGCAGTTCGAAAAATAA |
| iRFP702<br>with BsaI<br>overhangs | attaGGTCTCaCATGGCTCGTAAAGTCGACCTTACTTCCTGCGATCG<br>CGAGCCGATCCATATTCCGGGTTCTATCCAGCCATGCGGCTGCCTT<br>CTGGCTTGTGATGCTCAGGCCGTACGCATTACCCGTATTACGGAAA<br>ACGCCGGTGCTTTCTTTGGACGCGAAACACCTCGCGTAGGTGAAC<br>TGCTGGCCGACTACTTCGGCGAGACGGAAGCCCATGCCCTTCGTA<br>ATGCTCTGGCACAGTCGAGCGATCCAAAGCGTCCAGCCTTGATTT<br>TCGGATGGCGTGATGGCCTGACAGGGCGTACCTTCGATATTTCGTT<br>ACATCGTCACGATGGAACCTCGATTGTAGAGTTCGAGCCGGCGGC<br>GGCTGAACAAGCAGACAATCCTTTGCGTTTGACCCGCCAAATCATT<br>GCGCGCACGAAAGAACTTAAGTCGCTTGAGGAAATGGCGGCGCG<br>TGTGCCCCGTTATCTGCAAGCGATGTTAGGATACCACCGTGTCATG<br>CTTTATCGCTTCGCTGACGACGGGAGTGGCAAAGTAATTGGCGAG<br>GCCAAACGCAGCGATCTGGAGTCATTCTTGGGGCAACACTTCCCA<br>GCTTCCTTGGTTCCCCAACAAGCCCGCCTGTTATATCTGAAAAACG<br>CTATTCGCGTAGTTTCAGACAGTCGCGGTATTTCCTCTCGTATCGTA<br>CCCGAACATGATGCGTCGGGAGCCGCTTTGGACCTGAGCTTCGC<br>GCATCTGCGTTCTATCAGCCCGATTCATTTAGAATTTCTGCGTAATAT<br>GGGAGTTAGCGCATCCATGAGCCTTAGTATCATTATCGACGGAACT<br>TTGTGGGGCCTTATCATTTGCCATCACTACGAGCCCCGTGCGGTG<br>CCTATGGCTCAGCGCGTCGCTGCGGAAATGTTTGCAGATTTTCTTT<br>CGTTACACTTCACTGCTGCTCACCATCAACGCTAAAtGAGACCatta |
| mFAP2a<br>with BsaI<br>overhangs | attaGGTCTCaCATGTCACGCGCCGCGCAGCTCCTGCCGGGTACCT<br>GGCAGGCAACCTTTACCAACGAAGACGGTCAGACCTCCCAGGGC<br>CAGTGGCACTTCCAGCCGCGTTCTCCGTACACCATGGACATTGTT<br>GCTCAGGGTACCATCTCTGACGGTCGTCCGATCGTTGGTTACGGT<br>AAAGCGACCGTTAAAACCCCGGACACCCTGGACATAGACATCACC<br>TACCCGTCTCTGGGTAACATCAAAGCTCAGGGTCAGATAACCATGG<br>ACTCTCCGACCCAGTTCAAATGGGACGCAACCACCAAAGGTGAAA |

| | |
|---|---|
| | ACGATTTCCATGGTCGTCTGACCGGTACCCTGCAGCGTCAGGAAT<br>AAAtGAGACCatta |
| PFOR33 with Strep-Tag II | ATGGCTAGCTGGAGCCACCCGCAGTTCGAAAAAGCTAAAGAAAAG<br>AAGTTTATCACCTGTGATGGTAACGAGGCTGCAGCTCACGTGAGCT<br>ACATGTTCTCGGAGGTAGCTGCTATCTACCCCATCACCCCGTCTTC<br>GCCTATGGCGGAGCATGTTGACGAGTGGGCTGCCCGTGGTCGTAA<br>GAACCTGTGGGGCCAGAACGTCTCAGTTCAGGAAATGCAGTCAGA<br>GGCTGGTGCTGCCGGTGCCGTTCACGGTTCGCTGCAGGCTGGTG<br>CTCTCACCACTACATTCACCGCATCTCAGGGCTTGCTCCTGATGAT<br>TCCTAACATGTACAAGATTGCCGGTGAGCTGATTCCTTGCGTATTC<br>GACGTTTCTGCCCGTACGCTGGCTTCACACTCTCTGTGTATCTTTG<br>GCGACCACCAGGACGTAATGGCTTGCCGTCAGACCGGTTTTGCTA<br>TGTTCTGCTCAGGTTCAGTTCAGGAGGTTATGGACCTTACCGCTGT<br>TCCTCACCTGGCTACCCTCGAGACTATGGTTCCATTCGTAAACTTC<br>TTCGATGGTTTCCGTACCTCTCACGAGTATCACAAGATTGAGATGAT<br>GGACATGGAGGATATCCGTCCTCTGGTTAAGGAAGAGTACATCAAG<br>CGTTTCCGCGATCGTGCTATGAGCCCTGAGCGTCCTATCACTCGT<br>GGTACTGCCGAGAACCCCGAGACCTTCTTCACACACCGTGAGGCT<br>TGTAACCCATACTACGATTCAGTTCCCGAAGTAGTAGAGAAGTACCT<br>GGGCGAGATTTCAAAGATCACTGGTCGTGAGTACCACCTGTTCTC<br>ATACTACGGAGCCGAGGATGCCGACCGCATGATCATCCTCATGGG<br>TTCGGCTACCGATGCTGCACGTGAGGCTATCGACTACCTGAACGC<br>AAATGGTCAGAAGGTTGGTATGATCTCTGTTCACCTGTATCGTCCAT<br>TCTCAGTTAAGCACCTGCTGGCTGCTGTTCCTAAGACCGTTAAGAA<br>GATTGCCGTTCTGGATCGTACTAAGGAGCCAGGCGCTAACGGCGA<br>GCCTCTGTACCTCGACGTGAAGGATGCTTTCTACGATGTTCAGGAT<br>CGTCCTGTTATCGTTGGCGGTCGCTACGGTCTTGGTAGCCGCGAC<br>ACCACACCTGCTCAGATCATCAGCGTGTTCAACAACCTCGCTATGC<br>CCGAGCCCAAGAACCACTTCACCGTTGGTATCGTAGATGACGTCA<br>CCTTCACCTCTCTGCCTGAGGTTGAGGAGATTGCACTCGGTGGAA<br>AGGGCATGTTCCAGGCTAAGTTCTACGGTCTGGGTGCCGATGGTA<br>CCGTAGGTGCTAACAAGAACTCAGTTAAGATTATTGGCGACAACAC<br>CGACAAGTACTGCCAGGCTTACTTCTCTTACGACTCTAAGAAGTCG<br>GGAGGTTTCACCTGCTCTCACCTGCGTTTCGGTGATACACCTATCC<br>GCTCTACCTACCTGGTAACCACACCTAACTTCGTGGCTTGCCACGT<br>TCAGGCTTACCTCCACATGTACGACGTAACTCGTGGTCTGCAGAAG<br>AATGGTCAGTTCCTGCTGAACACCATCTTCGACGCCGACGAGCTC<br>GAGAAGTTCATGCCTAACAAGGTAAAGCGCTACTTCGCACAGAACA<br>ACATTACTGTTTATACCATCAACGCTACTAAGATTGCACAGGAGATT<br>GGTTTGGGCAACCGTACCAACACTATCCTGCAGAGTGCATTCTTCC<br>GCATCACCGGCGTTATCCCCGTTGAGCTCGCTGTTGAGAAGATGA<br>AGGCTGCCGCCCTGAAGTCTTACGGCGCTAAGGGTCAGGACGTT<br>GTTGATAAGAACTATGCCGCTATCGATCGTGGTGGTGAGTATGTTC<br>AGCTCACAGTTAAGCCTGAGTGGGCTAACCTGCCTGATGATGAGG<br>TTAAGGCCGACGACGCTCCCGCATTCGTTAAGGAGCTGGTTCGTC<br>CTATCAACGCTCAGGCCGGTGACCTGCTGAAGGTTTCTGACTTCG |

| | |
|---|---|
| | TTAAGCACAACACCGTTGATGGTTCTTGGGAGGTTGGTACAGCCG<br>CTTACGAGAAGCGTGGTGTTGAGGCCTTCGTTCCTGTATGGAACA<br>AGGACAACTGTATCCAGTGTAACCAGTGTGCTTACATCTGTCCTCA<br>CGCAGCTATCCGTCCATTCGTTCTCGACGACGAGGAACTGAAGGG<br>CTTCGCTGCTGCTGATGACACTATCGAGATGAAGGCACCTGCTGC<br>CATGAAGGGCATGCACTTCCGCATCGAGACATCTGTACTCGACTGT<br>CTGGGTTGCGGTAACTGTGCTGATATCTGCCCAGGCAATCCTAAGA<br>CTGGTAAGGCTCTCACCATGGCTCCATTCAATCCTGATGCTGCCGA<br>TATGGTTCAGGAGGCTAAGAACTGGGAGTACCTGGTTAAGAACGT<br>CAAGAGCAAGCAGAACCTCGTTGACATCAAGAGTAACGTTAAGAA<br>CTCACAGTTCGCTCAGCCTCTGTTCGAGTTCTCAGGCGCATGCTC<br>TGGTTGCGGTGAGACTCCATACGTTAAGCTGATTTCTCAGCTGTTT<br>GGTGATCGTCAGATGATTGCCAACGCTACTGGATGTTCTTCAATCT<br>ACTCTGCTTCTATTCCTTCTACACCTTATACCAAGAACGAGAAGGGT<br>CAGGGTCCTGTATTCAACAACTCTCTGTTCGAGGACTTCTGCGAGT<br>TCGGTCTGGGTATGGCTCTCGGTAACAAGAAGATGAAGGAGCGCG<br>TTGCTAAGCTGCTGGCTGAAGCTTGCGAGAAGTGCGACGTAACTC<br>CTGAGTACAAGGCTCTGGCCGAGGAGTGGATTGCCAACAAGGATG<br>ATGCCGACAAGACCAAGGAGATTGCTCCAAAGTTGCGCGCAGAGA<br>TTGCTGCTTGCGCCGCTAAGGGTTGCGAGATTTGCAAGGAGCTCC<br>AGACTCTGGATCACTACCTGGTTAAGCGCTCACAGTGGATCATCGG<br>TGGTGACGGTGCTTCTTACGATATCGGTTACGGTGGTCTCGACCAC<br>GTGATTGCTTCTGGCGAGGATGTAAACATCCTGGTACTCGATACCG<br>AAGTTTACTCTAACACCGGTGGTCAGGCTTCTAAGGCTACTCCTCT<br>TGGTGCTATCGCTCAGTTCGCTGCTCAGGGTAAGCGCATCCGCAA<br>GAAGGATCTGGGTATGATTGCCACAACCTACGGTTATGTATATGTAG<br>CTCAGATTGCTATGGGCGCCGACCACGCACAGACCCTCAAGGCCA<br>TCCGCGAGGCTGAGGCATGGCACGGACCTTCAATCATCATCGCCT<br>ATGCTCCATGTATCAACCACGGCTTGAAGGCCAAGGGCGGTATGG<br>GTAAGAGCCAGGCCGAGGAGAAGAAGGCTGTTGAGTGCGGTTAC<br>TGGCACCTGTGGCGTTTCAACCCAGCTCTGGCAGAGGAAGGCAA<br>GAACCCATTCTCACTCGACTCTAAGGAGCCCAACTGGGAGAACTT<br>CCACGACTTCCTGCTCGGCGAGGTTCGCTACCTGTCAGTTAAGAA<br>GGCTTATCCTAACGAGGCCGAAGAGCTCTTCGCTGAGGCTCAGAA<br>GATGGCCCAGCTCCGCTACAAGACTTACATCCGCAAGTCTCAGGA<br>AGATTGGACTGAGTAA |
